# Mathematical model for Plant-Insect interaction with dynamic response to PAD4-BIK1 interaction and effect of BIK1 inhibition

**DOI:** 10.1101/134619

**Authors:** Sanjay, Sabahuddin Ahmad, M. I. Siddiqi, Khalid Raza

## Abstract

Plant-insect interaction system has been a widely studied model of the ecosystem. Attempts have long been made to understand the numerical behaviour of this counter system and make improvements in it from initial simple analogy based approach with predator-prey model to the recently developed mathematical interpretation of plant-insect interaction including concept of plant immune interventions Caughley and Lawton (1981). In our current work, we propose an improvement in the model, based on molecular interactions behind plant defense mechanism and it’s effect on the plant growth and insect herbivory. Motivated from an interaction network of plant biomolecules given by Louis and Shah (2014) and extending the model of Chattopadhyay, et al (2001), we propose here a mathematical model to show how plant insect interaction system is governed by the molecular components inside. Insect infestation mediated induction of Botrytis Induced Kinase-1 (BIK-1) protein causes inhibition of Phyto Alexin Deficient-4 (PAD4) protein. Lowered PAD4, being responsible for initiating plant defense mechanism, results in degraded plant immune potential and thus causes loss of plant quality. We adapt these interactions in our model to show how they influence the plant insect interaction system and also to reveal how silencing BIK-1 may aid in enhanced production of plant biomass by increasing plant immunity mediated by increase in PAD4 and associated antixenotic effects. We hypothesize the significance of BIK-1 inhibition which could result in the improvement of the plant quality. We explain the interaction system in BIK-1 inhibition using mathematical model. Further, we adopted the plethora of computational modeling and simulations techniques to identify the mechanisms of molecular inhibition.

## Introduction

Insects are amongst the major threat to several economic and staple crops. Aphids make a significant bunch of contribution to this class of organism and annual crop loss^1^. Where insects have natural tendency to feed on plants and pose threat to their growth rate; plants also respond these herbivore attacks by initiating their defense mechanisms^2,3,4,5^. Altogether, these two form an ecological model of predator-prey system ^6^. We implement here the top-down approach of systems biology to further explain this model in mathematical expressions as well as to explain how molecular components behind this interaction helps or can be used for regulating this ecological system^7^. Though, a plethora of research works published till date have put forth the mathematical model of plant-insect interaction but none of those have yet included the molecular control responsible for this phenomenon^8,9,10,11,12,13^. In this work, we present a two-step workflow; of which one is to design a mathematical model of plant-insect interaction network in effect of molecular components behind their response; while other relates to the search of possible inhibitor against one of the key component of this network. At last we will returnto our mathematical model with an added component and show how this inhibition may affectnetwork and contribute towards protection of plants by synergizing itsdefense mechanism.

Some key research focused on elucidating the signaling mechanisms of plant defense systems has revealed molecular interactions involved in plant defense response^14,15,16,17,18,19, 20, 21^. A small module of such signaling mechanism have been reviewed and reported by Louis and Shah, (2014) ^22^. Proposed model is an extension to Plant-Insect interaction model supposed to analyze the synchronization between molecular level as well as community level change of phase of system components and their dynamic behavior. Moreover, we expect to analyze the change in behavior of our network’s dynamicity in effect of any perturbation in system at molecular level.

Our motivation to this work lies in studying trends of plant response on insect infestation causing rapid induction in PAD4 concentration. This increase in PAD4 concentration helps transcribing several other genes including those responsible for deterring insects and protecting plants from their attack ^21, 14, 23^. This response is accompanied by rise in Botrytis induced kinase (*BIK1*) protein concentration ^22^. BIK1 is a key molecule in plant defense system, as it seems to confer defense resistance against some insect herbivores in several plants. Phosphorylation of FLS2 and BAK1 proteins as well as association with Flg22 initiates BIK1 mediated defense signaling in such plants. As a final outcome of this signaling response, plants act through JA and SA mediated defense mechanism ^24, 25^. While in case of green peach aphid attack, BIK1 is induced by insect infestation and is an experimentally validated inhibitor of *PAD4* protein^14^.

Thus, in case of aphid attack sap feeding herbivores, insect infestation is not restricted because of *PAD4* inhibition in effect of *BIK1* expression. While, in *BIK1* mutants, basal activity of *PAD4* is up regulated and improves crop quality by hindering their infestation by insects^21^. This makes *BIK1* to be an important target of for inhibitor design against insect infestation of plants. Since, it seems quite difficult for any small molecule to cross cell wall barrier for entering inside cell and making it feasible for our inhibitor to serve this purpose. But we assume that the cell wall barrier inplants to be accessible by projected molecule upon leaf puncture during aphid attack and deliver its purpose in the cytoplasmic environment of plant. This will make a small molecular inhibition process of *BIK1* and may serve our purpose of improving plant growth rate through increased release of antixenotic phytochemicals, i.e. Ethylene ^15, 16^. Thus, in light of knowledge to this interaction and its anticipated effect on plant insect interaction system, Fig 1 shows a schematic diagram of the interaction network between Plants and Insects as well as their resultant influence on molecular components attached to it. We also try to demonstrate with the help of our mathematical model how inhibition of BIK1 can help restricting insect infestation and enhance plant biomass production.

**Figure 1:**
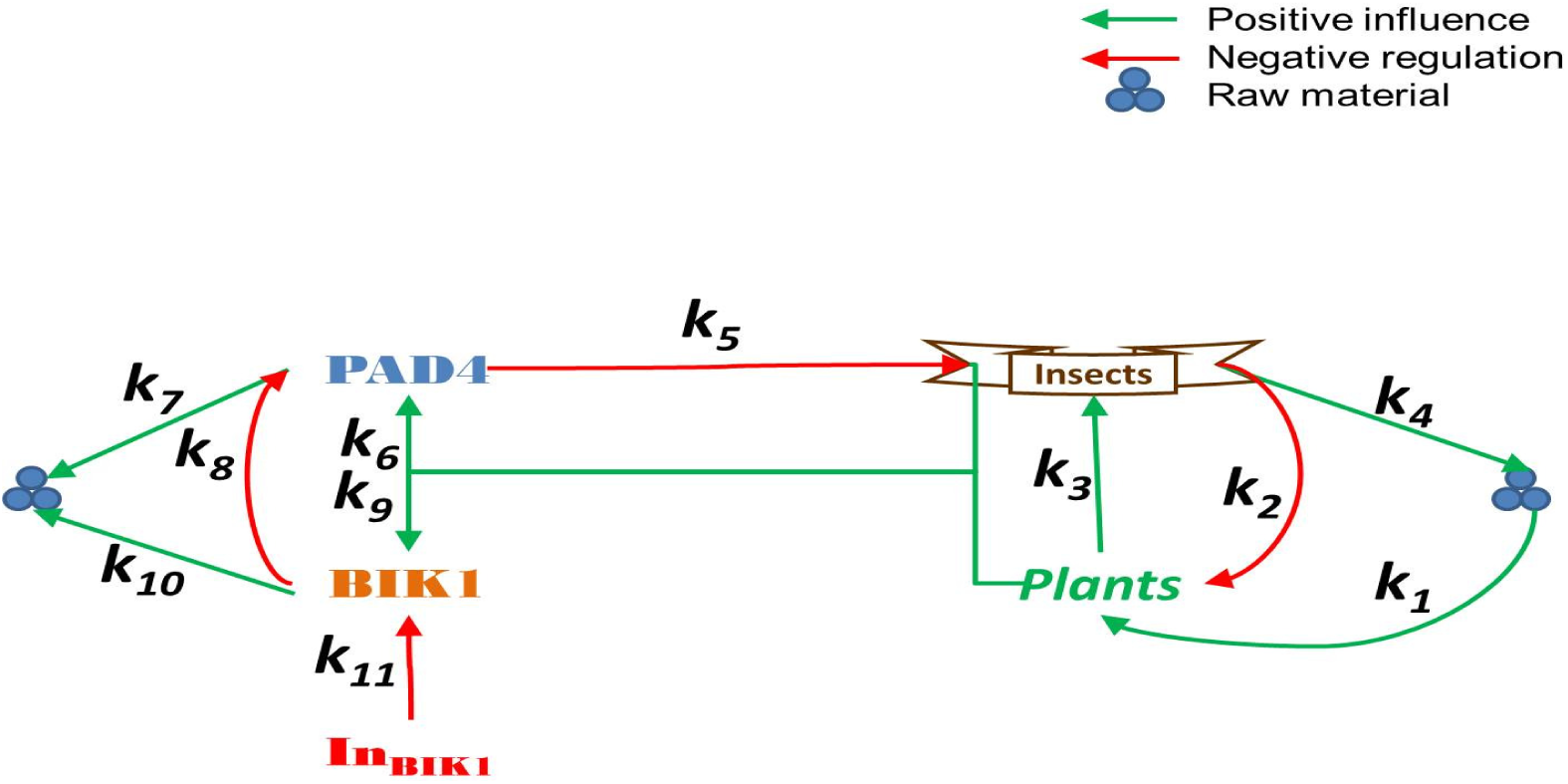
Schematic Diagram of proposed model.

## Methods

### Mathematical model

Efforts here in this work have been made to bring together Plant-Insect interaction model and its regulatory aspects in effect of molecular interactions involved. We restricted on basic behavioural interaction assuming sap feeding insects cause serious damage to plant stem and make it more difficult to recover as well as to simplify the computational complexity and focus on theme study of behavioral changes in effect of molecular interactions. This model consists of two parts, both of which showing dynamic progression of plant-insect interaction in effect of underlying BIK1-PAD4 interactions^22^; with an added inhibitor of BIK1 in the second step. Here, plants (***x***_***1***_) were considered as prey of growth rate ***k***_***1***_ as well as an assumption of long recovery time. Thus plants were considered to be damaged in any condition at a rate of ***k***_***2***_ in this predator-prey model. While, infestation of plants causeincrease in survival potency of insects (***x***_***2***_) and promotes reproduction with the factor of ***k***_***3***_. While reproduction of plant specieswas independent of any interaction and grew with the rate of *r* times its own population, but was limited by the carrying capacity of environment *K*. Since, there was no predator for insects in our model; they were limited by their own death rate of ***k***_***4***_. Further extension of this model was based on interaction of molecular components found during literature survey^10, 12^. Acording to which, we have introduced numerical dependencies of Plant-insect interaction system on underlying molecular interaction between PAD4 and BIK1.

Table 1 shows the list of components participating in this model. Rate constants of these interactions were decided to be arbitrary constants of such a range that does not makes any significant change to the model with or without these components. Parameters of added components in such range was seemed logical and justified because of our assumption that these molecular interactions already existed but not included while designing these plant-insect interaction models. Thus, literature based knowledge of antixenotic effects of PAD4 (***x***_***3***_) was translated in to this model with an effective rate of ***k***_***5***_ depending on its availability to insects^21, 23^. Synthesis of PAD4 was predicted to be at the rate of ***k***_***6***_ based on infestation caused by insects, while its natural degradation was supposed to be at the rate of ***k***_***7***_ of its own concentration^19, 20^. Since BIK1 (***x***_***4***_) had been responsible for inhibiting the actions of PAD4, this BIK1 based PAD4 inhibition was considered to be at the rate of ***k***_***8***_^14^.As suggested by literature survey, BIK1 was considered to be produced upon interaction of plants and insectsat therate of ***k***_***9***_^26, 27, 28^. While, it’s natural degradation rate was predicted to be ***k***_***10***_. List of all reaction channels and their rate of progression have been shown in Table 2.

**Table 1:**
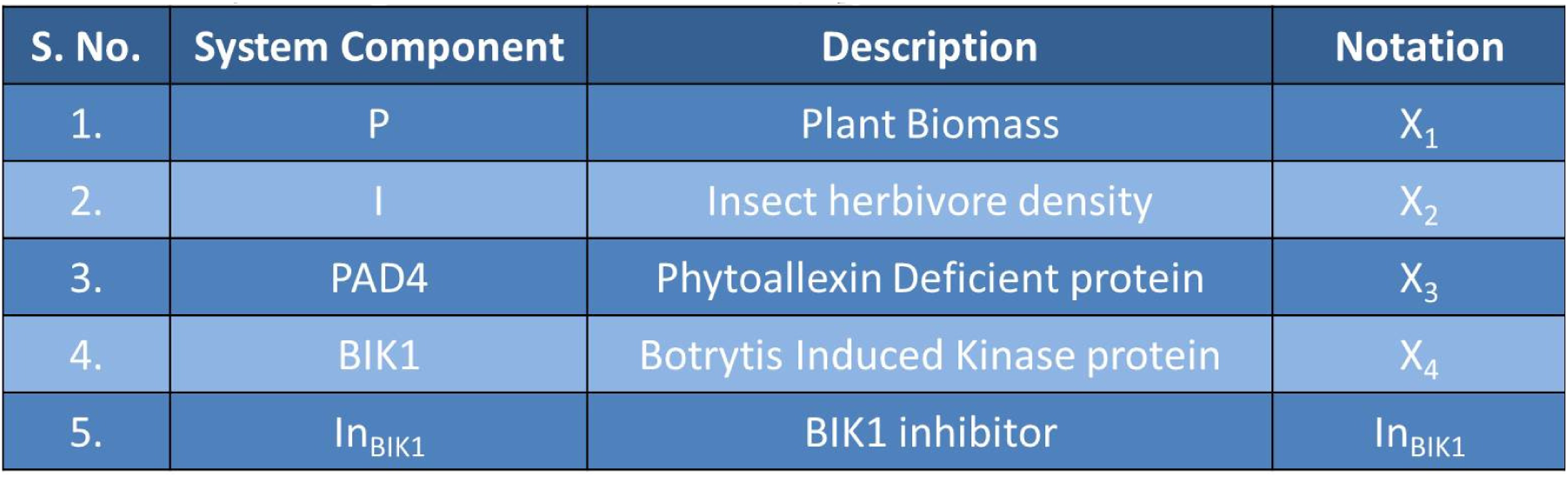
List of system components in extended network as proposed model.

**Table 2:** List of reactions involved in proposed system.

| S. No. | Biochemical reaction | Description | Rate Constants | Values of rate constants | Reference |
| --- | --- | --- | --- | --- | --- |
| 1. | $\phi \xrightarrow{k_1} x_1$ | Plant Biomass production | K1 | 0.2 | [10], [12] |
| 2. | $x_1 + x_2 \xrightarrow{k_2} x_2$ | Insect infestation on plants | K2 | 0.6 | [10], [12] |
| 3. | $x_1 + x_2 \xrightarrow{k_3} x_2 + x_2$ | Insect reproduction rate | K3 | 0.01 | [10], [12] |
| 4. | $x_2 \xrightarrow{k_4} \phi$ | Insect death rate | K4 | 0.02 | [10], [12] |
| 5. | $x_2 + x_3 \xrightarrow{k_5} x_3$ | Antixenosis by PAD4 | K5 | 0.002 | [21], [23] |
| 6. | $x_1 + x_2 \xrightarrow{k_6} x_1 + x_2 + x_3$ | PAD4 production | K6 | 1 | ----- |
| 7. | $x_3 \xrightarrow{k_7} \phi$ | PAD4 degradation | K7 | 0.1 | [19], [20] |
| 8. | $x_3 + x_4 \xrightarrow{k_8} x_4$ | BIK1 mediated PAD4 decrease | K8 | 0.1 | [14] |
| 9. | $x_1 + x_2 \xrightarrow{k_9} x_1 + x_2 + x_4$ | BIK1 production | K9 | 1 | [26], [27], [28] |
| 10. | $x_4 \xrightarrow{k_{10}} \phi$ | BIK1 degradation | K10 | 0.1 | ----- |
| 11. | $In_{BIK1} + x_4 \xrightarrow{k_{11}} In_{BIK1}$ | Inhibitor based BIK1 decrease | k11 | 0.1 | ----- |

With all the parameters and reaction channels in hand, mathematical model was translated into ODEs. System of equations **1**, **2**, **3** and **4a** holds quality of an intact system.

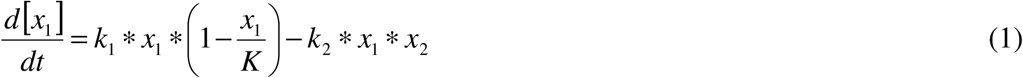

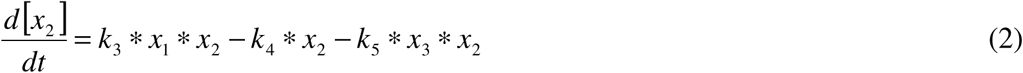

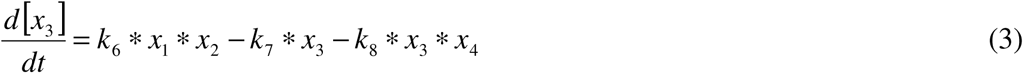

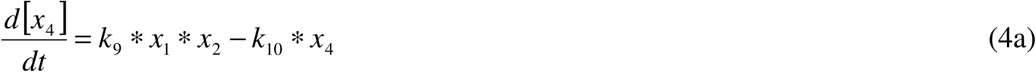

We used Runge-Kutta’s fourth order differential equation solver and PERL programming for simulation of this model^29^. It was made sure through model simulation, that no significant difference is being observed in network progression while extending the network with added PAD4 and BIK1.

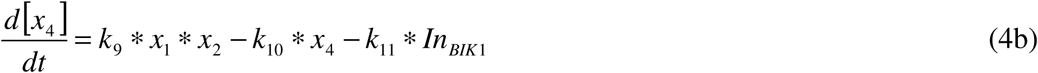

For, our purpose of studying effect of BIK1 inhibition on Plant insect interaction, we deliberately introduced a hypothetical inhibitor (***In***_**BIK1**_)in the model working at an effective rate of ***k***_***11***_. This change in network has been incorporated in the form of equation **4a** replaced by equation **4b** in our earlier system of ODEs. Simulation of this model is supposed to show a prediction of effect exerted by plants on the insect herbivores through enhanced antixenotic properties or more appropriately, an enhanced release of Ethylene through the plants^15, 16^. While, we fixed the minimum value limit of BIK1 decline to zero concentration for avoiding any negative value if achieved.

### BIK1 structure prediction and inhibition

As suggested by our mathematical model and experimental evidence the importance of BIK1 inhibition resulting in the higher PAD4 concentration in plant species thus inhibiting the aphid attack on plants. Based on the computational modeling and molecular dynamics simulation approaches we identify possible lead molecules, which can act as inhibitors to BIK1 in plants^30, 31^. Due to unavailable crystal structure, we decided to choose *ab intio* structure modeling through Phyre2 web server and minimized structural energy through FG-MD webserver. There onwards, we tested our protein structure for structural validation through Ramachandran plot analysis of Phi-Psi angle allowance as well as ProSA analysis for structural comparison with other PDB based structures. Emphasis structural validation was made to reach closest comparison of its crystal structure result.

### Sequence information and structure modeling

We retrieved the intact BIK1 protein sequence of *Arabidopsis thaliana* (NP_181496.1, 395 residues) from NCBI-Protein databank resource (https://www.ncbi.nlm.nih.gov/protein/). Figure 2a shows the information of GOR4 detected secondary structures available in sequence as well as active site region of the protein sequence detected from PSI-BLAST are shown in Fig 2b. GOR4 analysis predicted that BIK1 protein sequence consist of 137 amino acids i.e. 34.68% protein in alpha helix form, 62 amino acids i.e. 15.70% protein in extended strand form and random coil of 196 i.e 49.62% amino acid residues ^32^.

**Figure 2:**
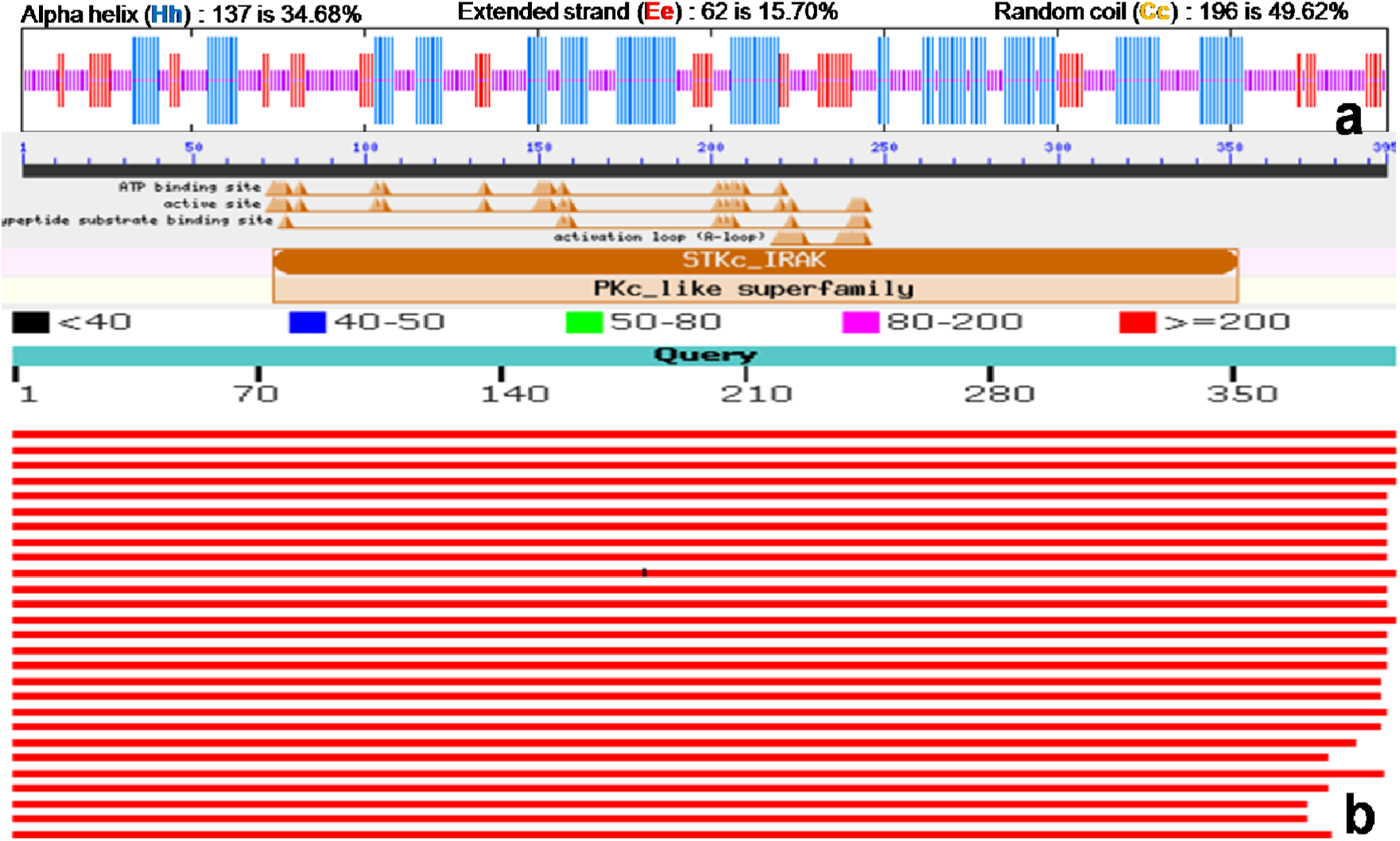
**(a)** Secondary structure analysis of BIK1 protein sequence through GOR4 confirming 34.68% alpha helix, 15.70% extended strand and 49.62% of random coil composition in the whole protein sequence 32. **(b)** PSI-BLAST result shows the putative range of active site, peptide substrate binding site and ATP binding site in the protein sequence ^33^.

PDB directed PSI-BLAST results were obtained to detect the feasibility of this protein’s homology modeling ^34^. Below 40% similarity index of BIK1 protein sequence with any crystal structure persuded us to build our protein’s structure through *ab initio* structure modeling techniques ^35, 36, 37^. Moreover, low similarity of this protein was confirmed from any of proteins in humans. This also strengthens our idea of using inhibitor of this protein during agricultural practices, without posing any threat to humans upon consumption in small amounts. We submitted our sequence to Phyre2 webserver for *ab initio* structure modeling at Intensive mode of setting. Shown below in Fig 3 is the structure obtained with 90% accuracy score and 80% query coverage ^38^.

**Figure 3:**
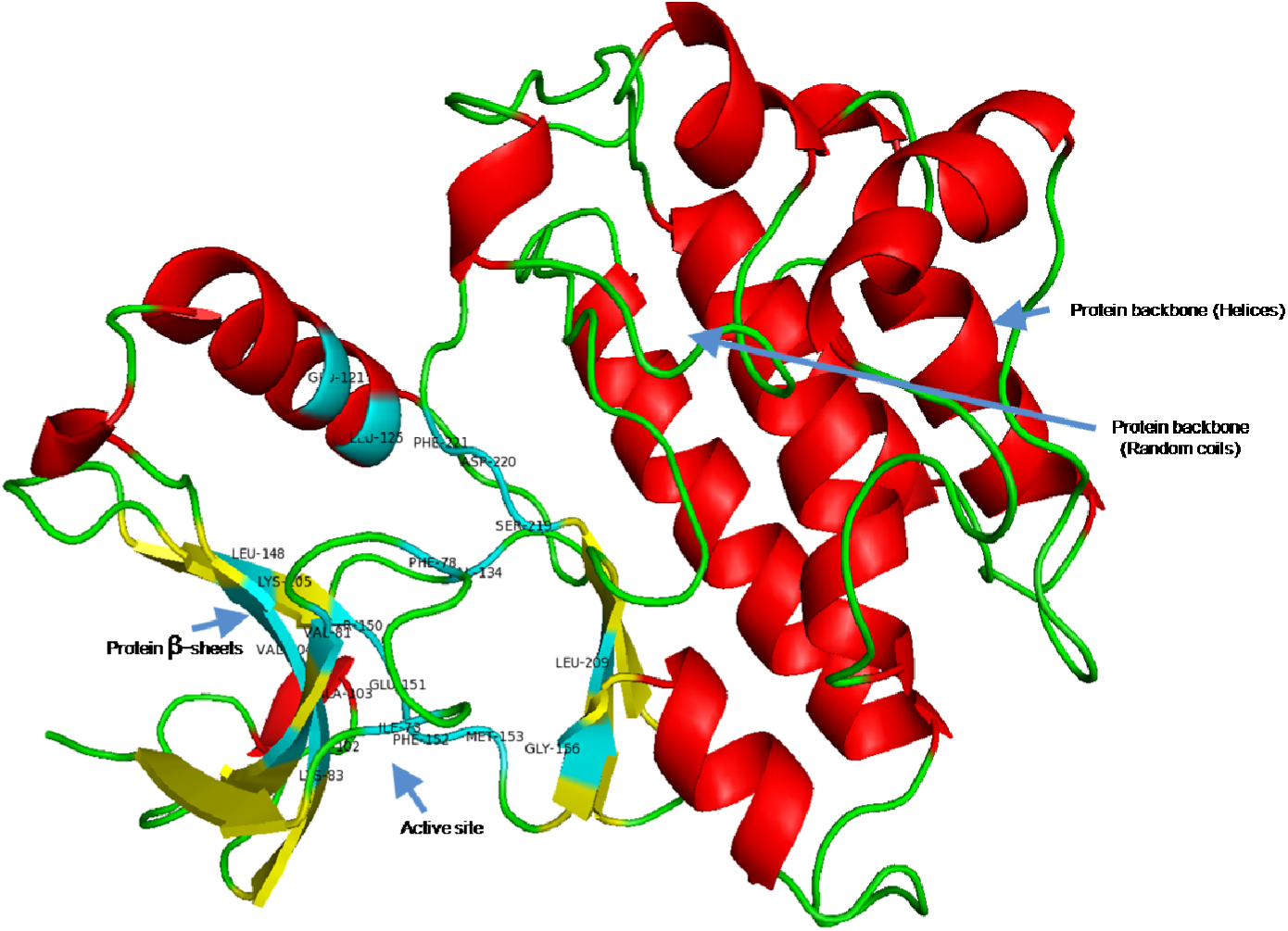
Structure of BIK1 protein modeled through Phyre2 server with 90% accuracy and 80% query coverage and calibrated for whole structure enrgy minimization through I-TASSER’s FG-MD server ^38, 39^.

At this stage, obtained PDB file was subjected to Fragment Guided Molecular Dynamic simulation server of I-TASSER for minimization of energy in PDB structure and get a structure with most stable conformation of predicted model ^39^. With this structure, we used RAMPAGE server Ramachandran analysis for structure validation confirming occurrence of 89.7% residues in favored region and 8.1% residues in allowed region (Fig 4a) ^40^. We also processed our protein through ProSA-web server to calculate the Z-score of this structure and Knowledge Based Energy diagram for the amino acid residues in this structure (Fig 4b & 4c) ^41^. Z-score for this protein structure was obtained -6.79 amongst all available structures in RCSB-PDB.

**Figure 4:**
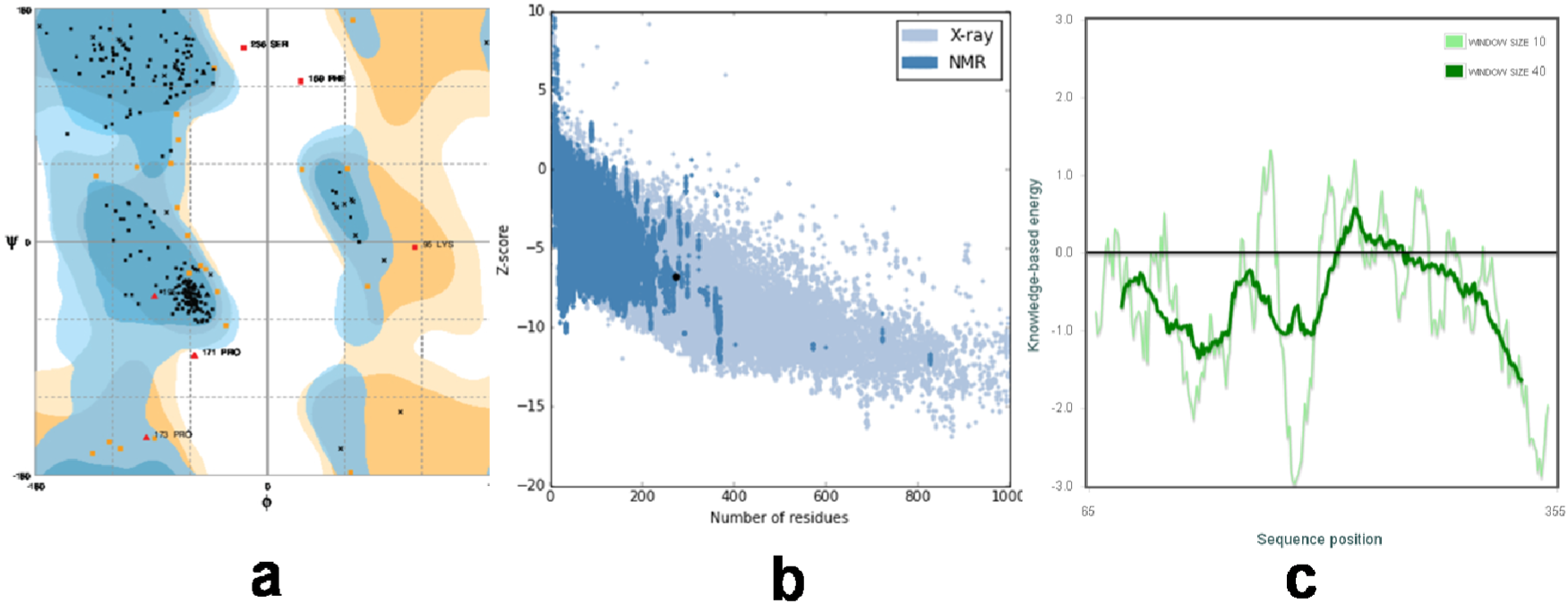
Protein validation report. **(a)** Ramachandran plot shows expected percentage of ∼90% residues in favoured region and ∼8% residues in allowed region; only <2% residues were observed as outlier amino acids ^40^. **(b)** Z-score plot of our predicted and Fragment Guided MD based refined model showing value of -6.79 in comparison to available structures in the RCSB-PDB database ^39, 41^. **(c)** ProSA-web based study of protein structure shows the residuewise energy of amino acids ^41^.

Thus, validating the all structural aspects of this model at *in silico* platform, we moved ahead in direction finding an inhibitor for this protein.

### Active site prediction

This protein model was then analyzed through METAPOCKET server as well as DogSiteScorer web server for studying surface topology of BIK1 protein structure and finding probable active sites ^42, 43, 44^. Table 3 shown below is an extension of active site region analysis through CD-search database and confirms the active region of BIK1 protein through results obtained from Pfam, Conserved Domain search, TIGR, PLN, COG and SMART motif search results^45^. We have summarized the participating amino acid residues in the active region of this protein in accordance with results obtained from PSI-BLAST. We also correlated PSI-BLAST confirmed active site region of this protein with surface topology result of DogSiteScorer server and submitted our protein for METAPOCKET analysis^43, 33, 46^. Correlation of these two analysis confirmed P_2 from DogSiteScorer analysis as the active and binding region of this protein.

**Table 3:**
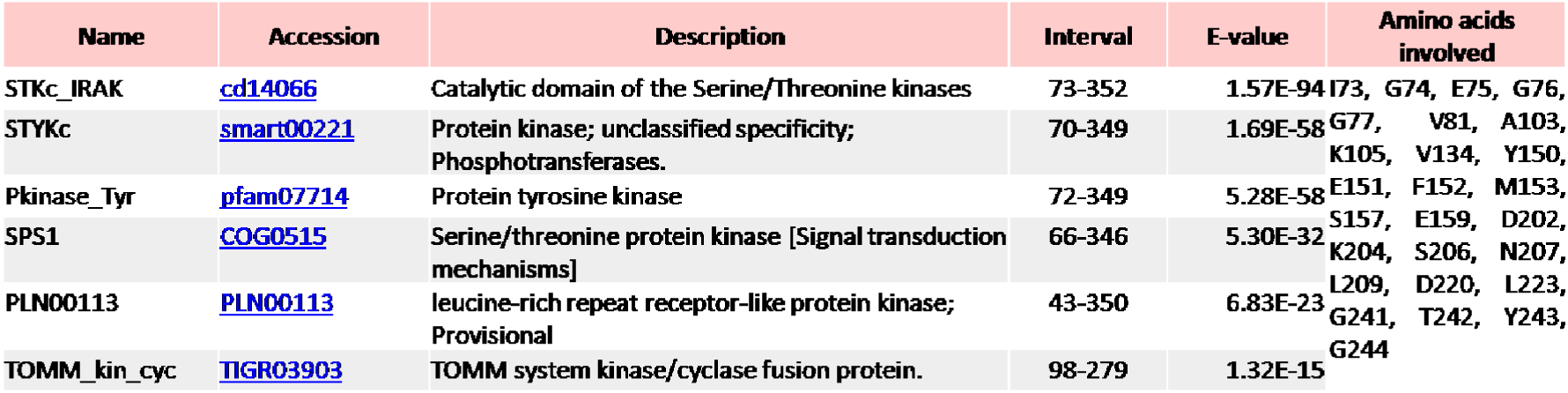
Conserved Domain analysis of BIK1 protein sequence showing putative range of active/binding domains retrieved from information available on different reliable databases **^45^**, **^47^**, **^48^**.

Our results from METAPOCKET detected same location with exactly same amino acids as Pocket 1 and strengthened our knowledge of the active site residues Table 4 shows a summary of DogSiteScorer detected pockets at the surface of BIK1 protein with details.

**Table 4:** Details of pockets available on the surface of BIK1 detected from DogSiteScorer with participant amino acids and their count **^43^**.

| Pocket_ID | Volume[Å<sup>3</sup>] | Surface[Å<sup>2</sup>] | SimpleScore | DrugScore | Amino acids involved (for P2 as active site only) |
| --- | --- | --- | --- | --- | --- |
| P_0 | 790.14 | 1297.4 | 0.54 | 0.818297 | **1 Alanine, 1 Aspartate,**<br>**2 Glutamine, 1 Glycine,**<br>**2 Isoleucine, 3 Leucine,**<br>**2 Lysine, 1 Methionine,**<br>**3 Phenylalanine, 1 Serine,**<br>**1 Tyrosine, 3 Valine** |
| P_1 | 590.98 | 937.38 | 0.4 | 0.760281 |  |
| P_2 | 407.94 | 451.45 | 0.26 | 0.732549 |  |
| P_3 | 395.46 | 681.26 | 0.17 | 0.734475 |  |
| P_4 | 354.24 | 679.72 | 0.19 | 0.568768 |  |
| P_5 | 280.58 | 266.71 | 0 | 0.755472 |  |
| P_6 | 236.22 | 462.54 | 0.07 | 0.424016 |  |
| P_7 | 194.37 | 383.76 | 0.03 | 0.428296 |  |
| P_8 | 130.24 | 207.99 | 0 | 0.335201 |  |
| P_9 | 114.18 | 232.82 | 0 | 0.243989 |  |

Comparing the results of DogSiteScorer, METAPOCKET and PSI-BLAST based study as well as manually detecting the location of participating amino acids, we concluded P_2 as the active site.

Figure 5 shown above is a structural representation of P_2 pocket of BIK1 structure specificying active site residues.

**Figure 5:**
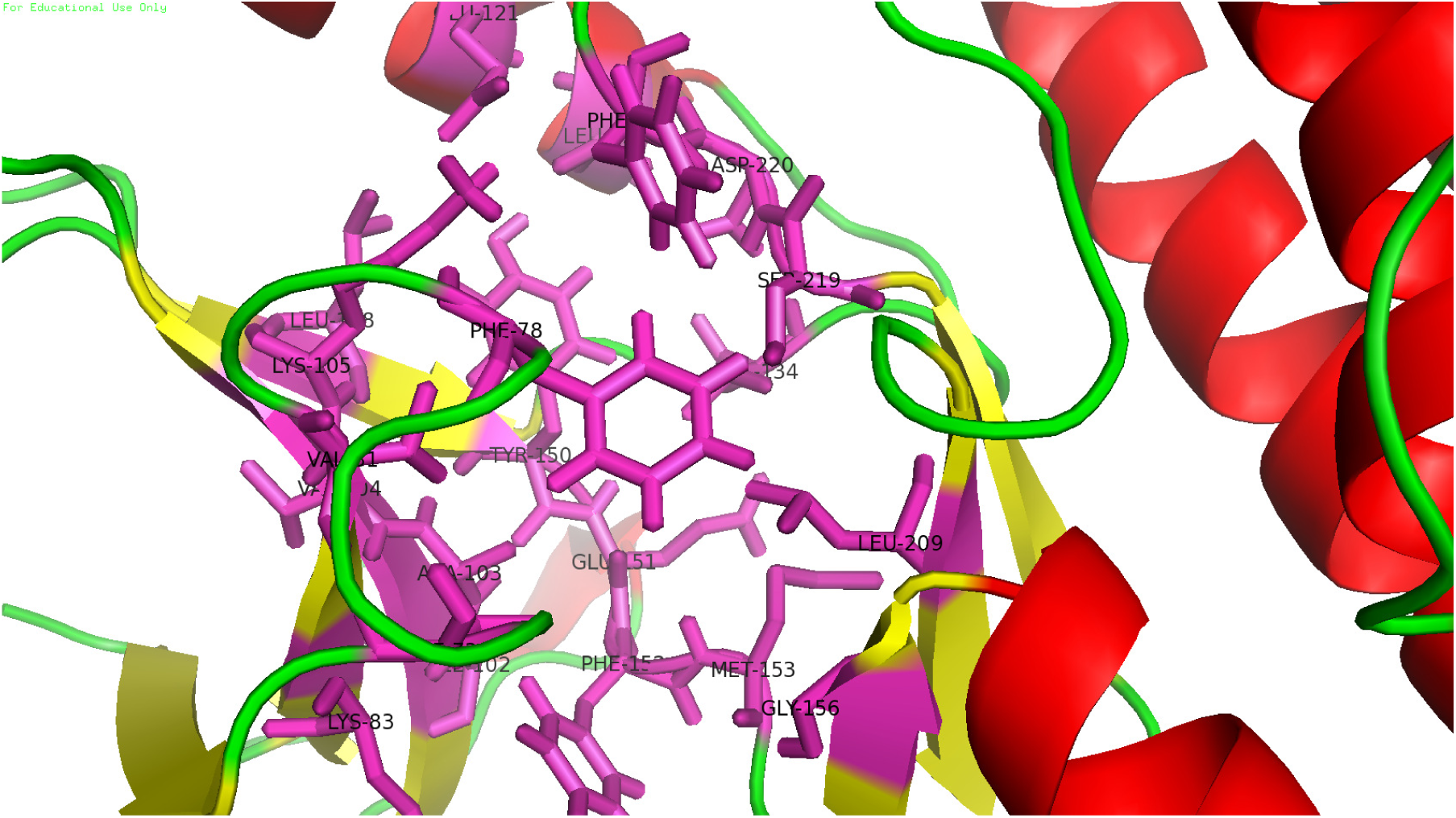
Surface pocket showing active site residues of the BIK1 protein structure obtained from DogSiteScorer and METAPOCKET analysis and confirmed from PSI-BLAST and CD-search results ^45, 33, 43, 44^.

### Protein preparation

The modelled structure of BIK-1 was subjected to Protein Preparation Wizard utility in Maestro (Schrödinger, Inc.). Hydrogen atoms were added to the resulting structure, which was minimised to overcome bad steric contacts. During receptor grid preparation wizard, OPLS/AA charges^49^ were automatically assigned by Glide version 7.1 (Schrödinger, Inc.).

### Ligand preparation

Structure files of Maybridge screening collection were obtained from its website (http://www.maybridge.com). There are approximately 53,000 organic molecules available at this time for screening purpose. Considering the LigPrep program (Schrödinger, Inc.), possible tautomers were generated using ionization states at pH 7. OPLS ligand charges were also assigned by the program to these structures at default setting.

### Docking studies

Docking calculations were completed in three stages. First stage included docking of all the possible tautomers of the available molecules of Maybridge library using High-Throughput Virtual Screening (HTVS) mode. The molecules which had Glide Score (GScore) below -6.0 were selected for next stage docking. In the second stage, Standard Precision (SP) mode was considered. Those molecules which had GScore below -8.0 were considered for final stage of docking. This stage included molecular docking using Extra-Precision (XP) mode of glide. The results were analysed and interpreted on the basis of visual inspection of molecular interactions with the protein using the Maestro (Schrödinger, Inc.).

### Molecular Dynamics

Molecular Dynamics (MD) simulations were carried out to validate the docking results. For this simulation, we have considered the GROningenMAchine for Chemical Simulations^50^ (GROMACS) v5.0.7 package. Topology file for the protein was prepared considering ‘GROMOS96 43a1’ force field^51^ and ligand topology was prepared over PRODRG server^52^. The solvated system was found to have a net charge of -2.0. Therefore, two sodium ions were added to neutralize the system. In next stage, solvated, neutral system was minimized using the steepest descent method, followed by position restraint dynamics, also known as equilibration run inclusive of NVT (Number of particles, Volume and Temperature) and NPT (Number of particles, Pressure and Temperature). Soon after the system was equilibrated, a 20 nanoseconds (ns) long production simulation (MD run) was initiated. The output files of the MD run were processed for analysis of the simulation performed using GROMACS. To interpret the hydrogen bonds occupancy for specific residues of BIK-1, VMD^53^ was used.

## Results and Discussion

### Mathematical model

We simulated our model on numerical basis and demonstrated how molecular responses during insect infestation of plants can be utilized in favor of human benefits by increasing plant growth rate by cessation of insect infestation. Fig 6a shows a damping oscilation of plant growth in normal conditions of insect infestation. In this plot, plant biomass accumulation stabilizes at 2.2533 concentration scale of this system in comparison to 105 carrying capacity supported in this model, while insect population stabilizes to the value of 0.3256. Thus, this system at ecological level became stationary by the end of 2507^th^ time scale at about two percent or less of the total carrying capacity of the ecosystem for plants. Meanwhile, molecular concentration of PAD4 and BIK1 also proceed through an oscilatory response as shown in Fig 6b. By the end of 2525^th^ time scale, both of these molecular components became stable at 0.8811 for PAD4 and 7.3318 for BIK1. These two plots can be easily correlated for PAD4 dependent insect population as well as respective plant quality. Fig 6c shows how relative amount of plant biomass vary in response to insect population. As soon as insect population increases, plant biomass dives down its relative amount. While, molecular dependency of plant biomass and insect population can be correlated with BIK1 and PAD4 concentration inside cell as seen in Fig 6d. As soon as BIK1 concentration rises, it drops down the relative concentration of PAD4 protein inside cell; this in response helps insects through lowered PAD4 based antixenotic effect and negatively affects relative plant biomass in the system.

**Figure 6:**
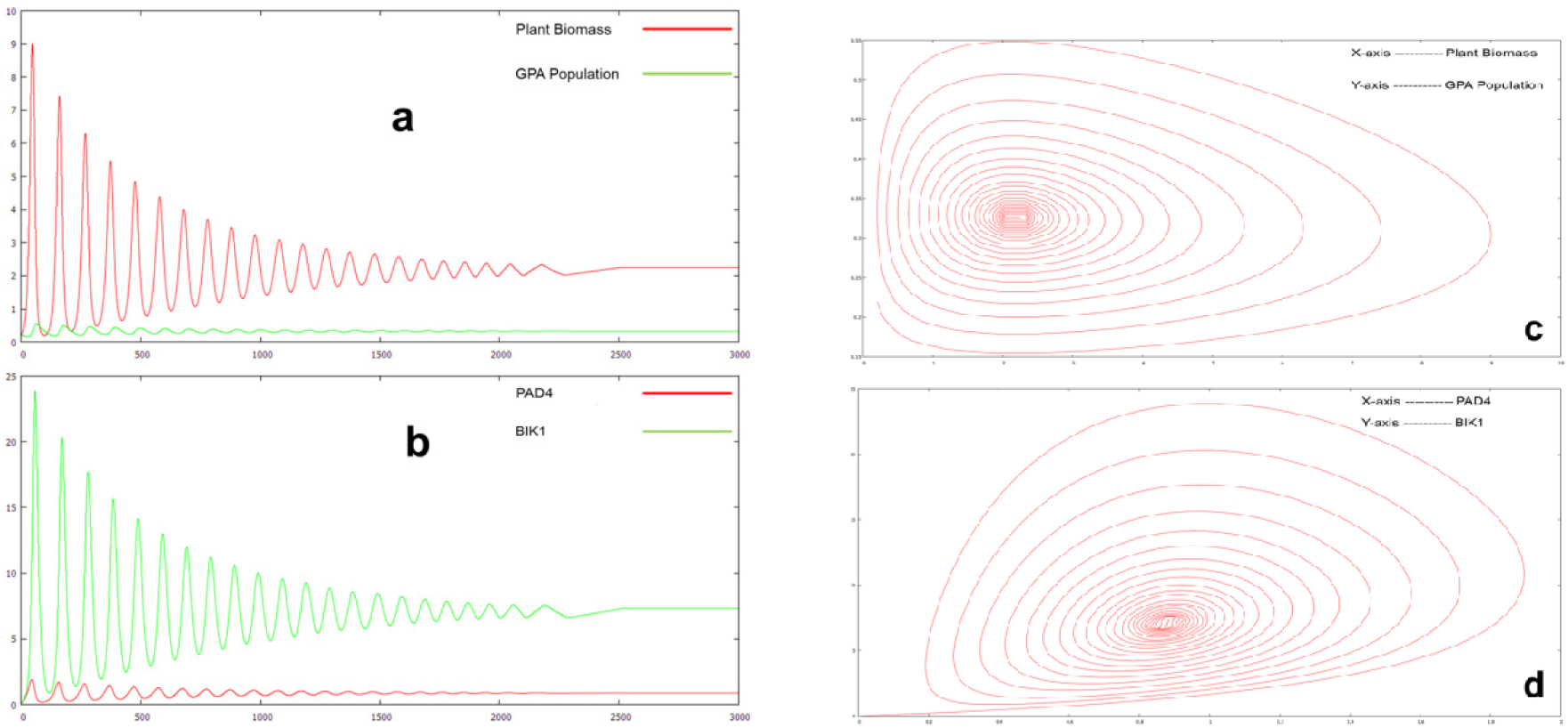
**(a)** Variation of Plant Biomass accumulation and GPA population in the system during regular insect attack without any pesticide or infestation inhibitor. **(b)** Underlying progression of PAD4 and BIK1 concentration at cellular level. **(c)** Relative dependency of plant biomass and GPA population at ecological system level response. **(d)** Concentration plot of PAD4 and BIK1 showing their relative dependecy on each other.

Fig 7 shown below summarizes the variation in system behaviour in response to introduction of BIK1-inhibitor at about 310^th^ time scale till next 20 time steps at the rate of 0.1 concentration scale per time step. We observed in Fig 7b, that BIK1 inhibition significantly increased the PAD4 concentration in plants and helped them deteriorating the insects to cease on their feeding preference.

**Figure 7:**
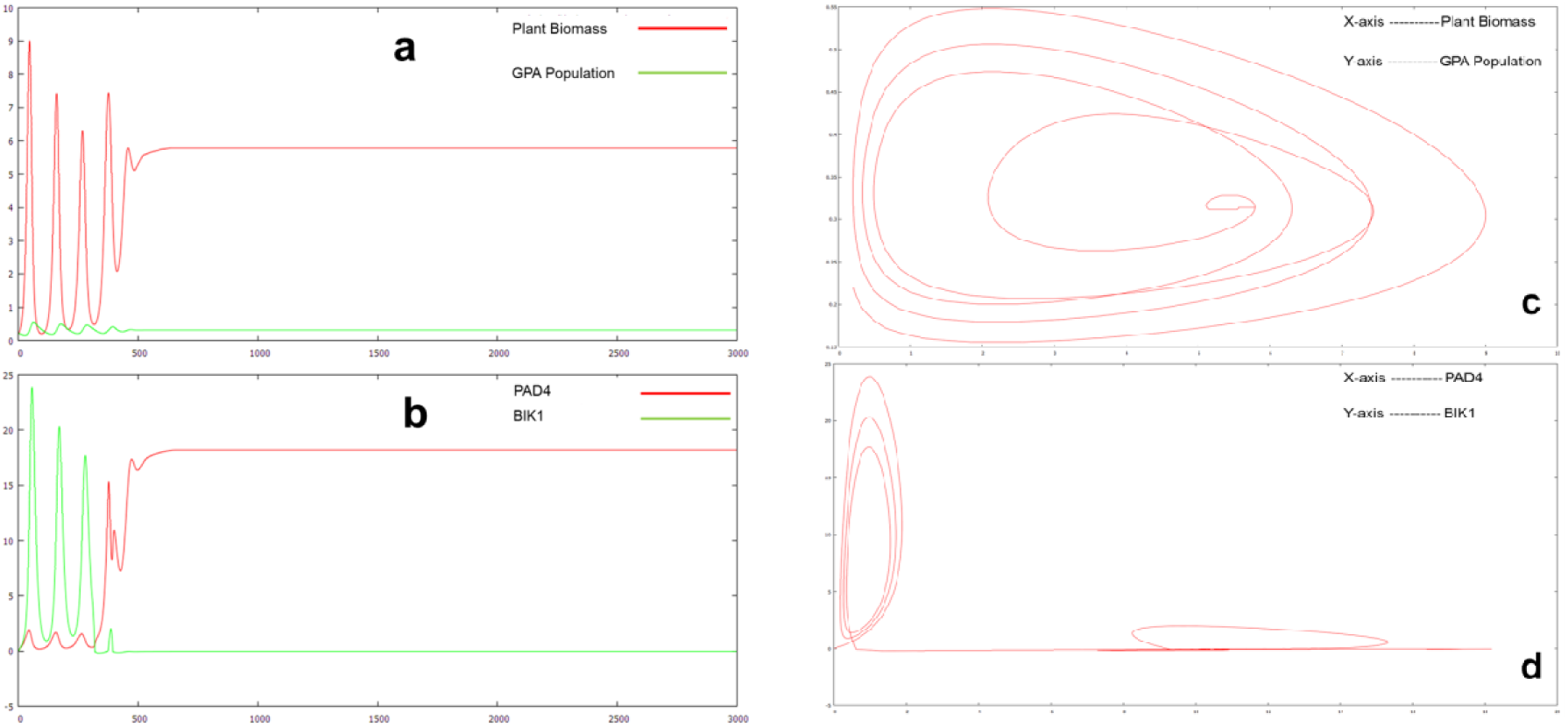
**(a)** Plot showing BIK1 inhibition based change in variation of Plant biomass and GPA population engaged in infestation from 310th time scale. **(b)** Plot indicating the sudden increase of PAD4 concentration at cellular level right after introduction of BIK1 inhibitor in the system. **(c)** Relative variation of Plant Biomass and GPA population in the system showing their dependency on each other upon addition of BIK1 inhibitor. **(d)** Plotted results of underlying PAD4-BIK1 concentration relative to each other and their response in addition to inhibitor against BIK1.

This inhibition resultantly helped keeping insects away from plants and increasing their biomass production as shown in Fig 7a. Thus, according to Fig 7d molecular control of BIK1 action directly influenced the action of PAD4 protein in the system, hence helped plant biomass growth rate to increase and get stable at about 5.7886 biomass scale by the end of 634^th^ time scale in contrast to 2.2533 during regular infection scenario. Meanwhile, by the end of 656^th^ time scale, BIK1 concentration reached zero level of concentration and stabilized PAD4 at 18.1944. Thus, according to numerical simulation and their result analysis, we concluded BIK1 inhibiton as the key mechanism to increase the plant growth. We, furthermore suggest that the numerical simulation may not have been able to completely mimic the real scenario and have only partly symbolized the effect of this inhibition. But, it might have a more influencial effect on the ecological system as some of the wet lab results showing effects of BIK1 silencing have indicated more promising plant growth.

### Virtual Screening and Molecular Docking

Predicted binding site of BIK-1 was considered for glide grid preparation and has been targeted for docking of molecules. All the possible tautomers of ∼53, 000 molecules were docked using HTVS mode of Glide. Around 23, 000 molecules had glide score equal to or below -6.00. These molecules were considered for molecular docking via SP mode of glide. There were around 1300 molecules which had glide score -8.0 or less. These molecules were further docked using XP mode of glide. HTVS and SP mode of glide offers molecular docking at lower computational costs than XP mode of glide. XP mode of glide is an important method as it has the ability to weed out the false positives at the same time providing a better correlation between both the good poses as well as good scores. From the XP glide docking results, the top 86 molecules having glide score less than or equal to -10.0 were short-listed for detailed analysis. Visual inspection of the docked complex was carried out. Based on maximum number of interactions of ligand (hydrogen bond, hydrophobic interactions etc) with the amino acid residues constituting the binding site the list was ranked. The top two ligands (i.e. RF03436 and SPB07243) had maximum interactions with the binding site of the protein (Table 5). Ligand interaction diagram in maestro provides vital information about the type of interactions between the ligands in complex with proteins.

**Table 5:**
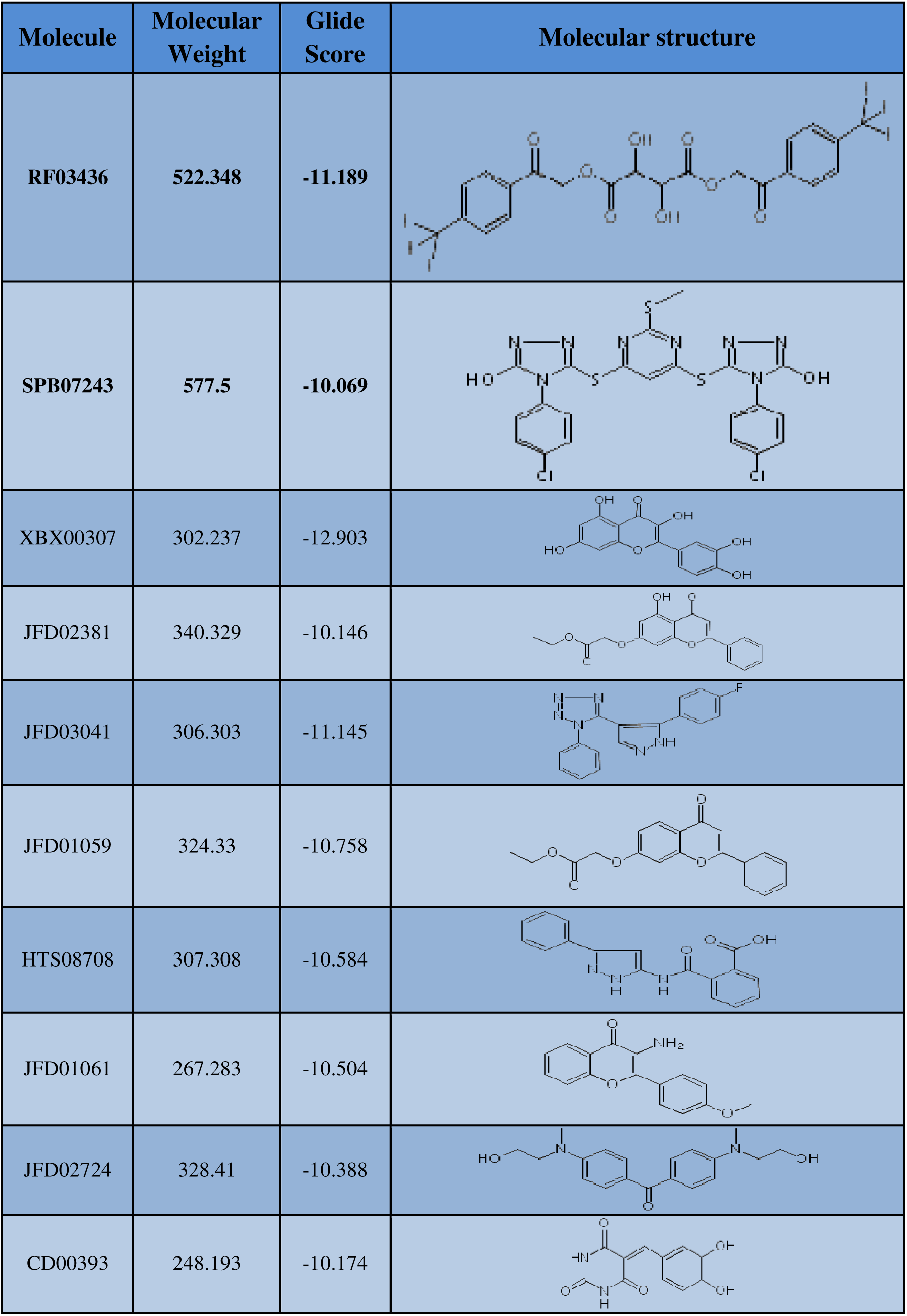
Molecular Docking results for top 10 molecules available from XP mode of Glide.

| Molecule | Molecular Weight | Glide Score | Molecular structure |
| --- | --- | --- | --- |
| RF03436 | 522.348 | -11.189 |  |
| SPB07243 | 577.5 | -10.069 |  |
| XBX00307 | 302.237 | -12.903 |  |
| JFD02381 | 340.329 | -10.146 |  |
| JFD03041 | 306.303 | -11.145 |  |
| JFD01059 | 324.33 | -10.758 |  |
| HTS08708 | 307.308 | -10.584 |  |
| JFD01061 | 267.283 | -10.504 |  |
| JFD02724 | 328.41 | -10.388 |  |
| CD00393 | 248.193 | -10.174 |  |

### Molecular interactions

Ligand interaction diagram of RF03436 (Fig 8) reveals that the molecule interacts with the protein via hydrogen bonds with amino acid residues constituting the binding site of the protein (i.e. Gly74, Met153, Ser157, Asn160). Moreover, there is also a pi-cation interaction with Arg164, a binding site residue. Several residues of the protein were found to form hydrophobic contacts with the RF03436 (Fig 8).

**Figure 8:**
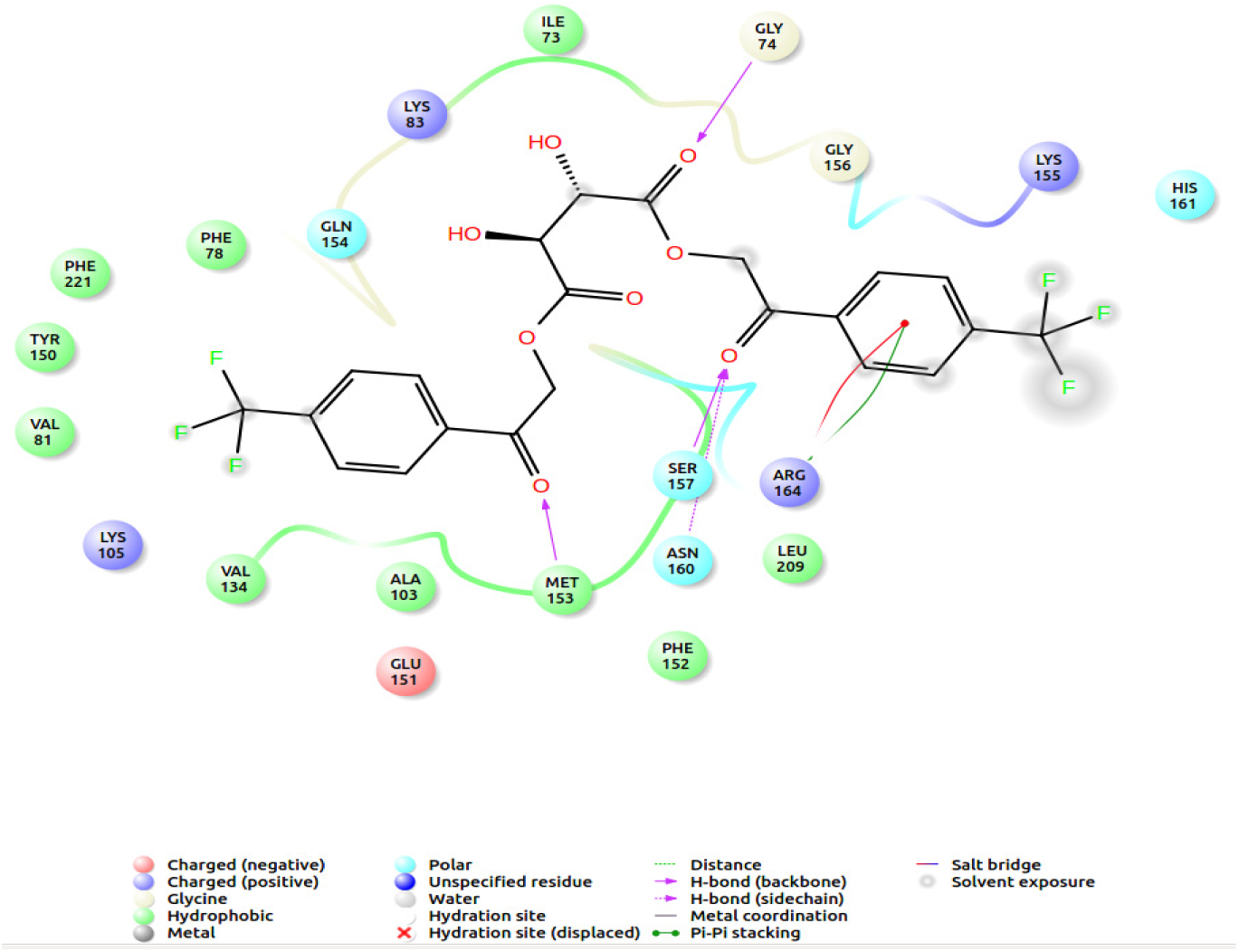
Ligand interaction diagram: RF03436 and modeled BIK-1.

Another view of the docked complex (Fig 9) reveals the position of the docked RF03436 within the binding site of BIK-1.

**Figure 9:**
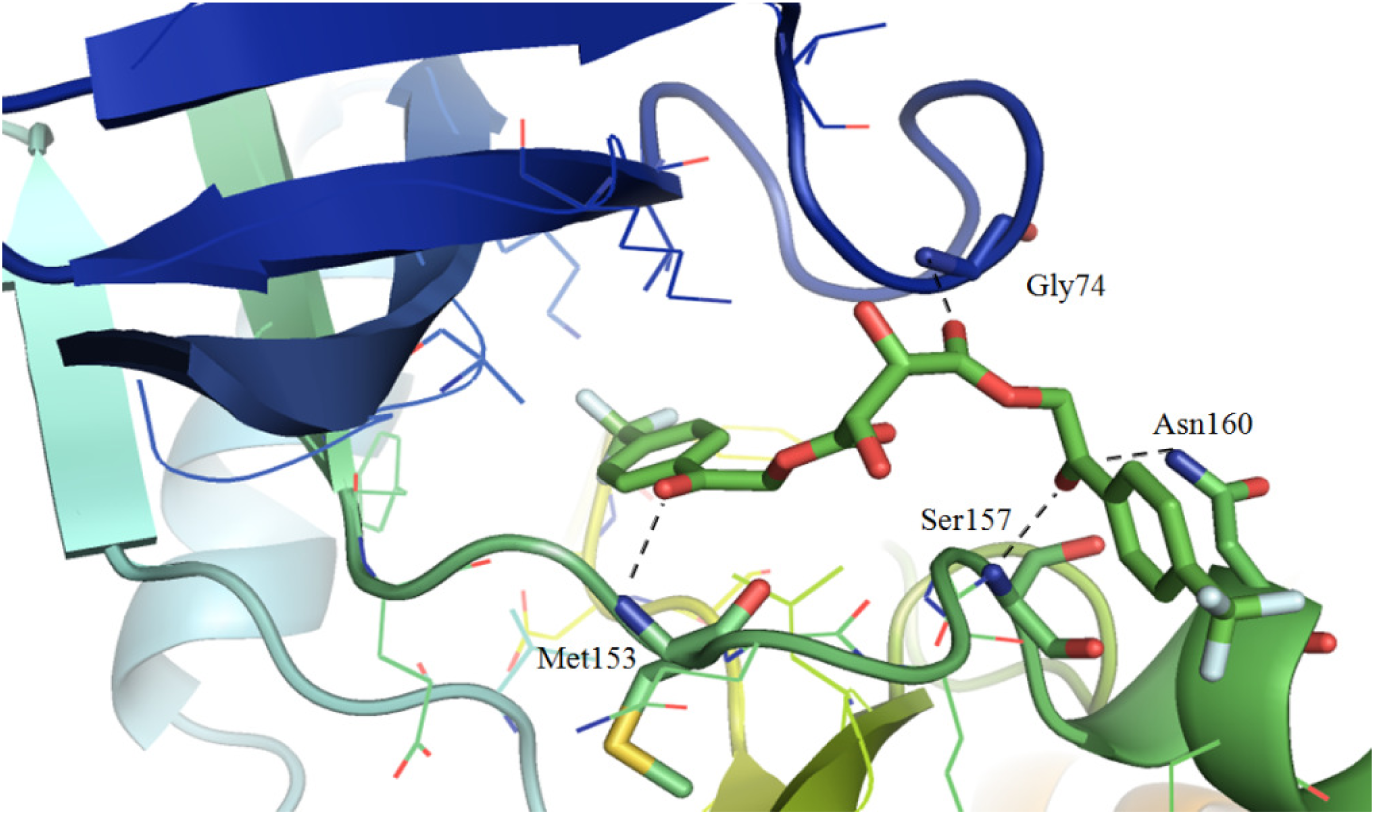
Comprehensive detail of interactions between BIK-1 and RF03436. Black lines represent hydrogen bonds.

For SPB07243, the ligand interaction diagram (Fig 10) describes that the molecule is making hydrogen bonds with various binding site residues of the protein like Val72, Gly74 and Met153. In addition, the ligand is also making pi-pi interaction with Phe78 and pi-cation interaction with Lys83, which are also the binding site residue of the protein. Several hydrophobic interactions are also involved between SPB07243 and the target protein (Fig 10).

**Figure 10:**
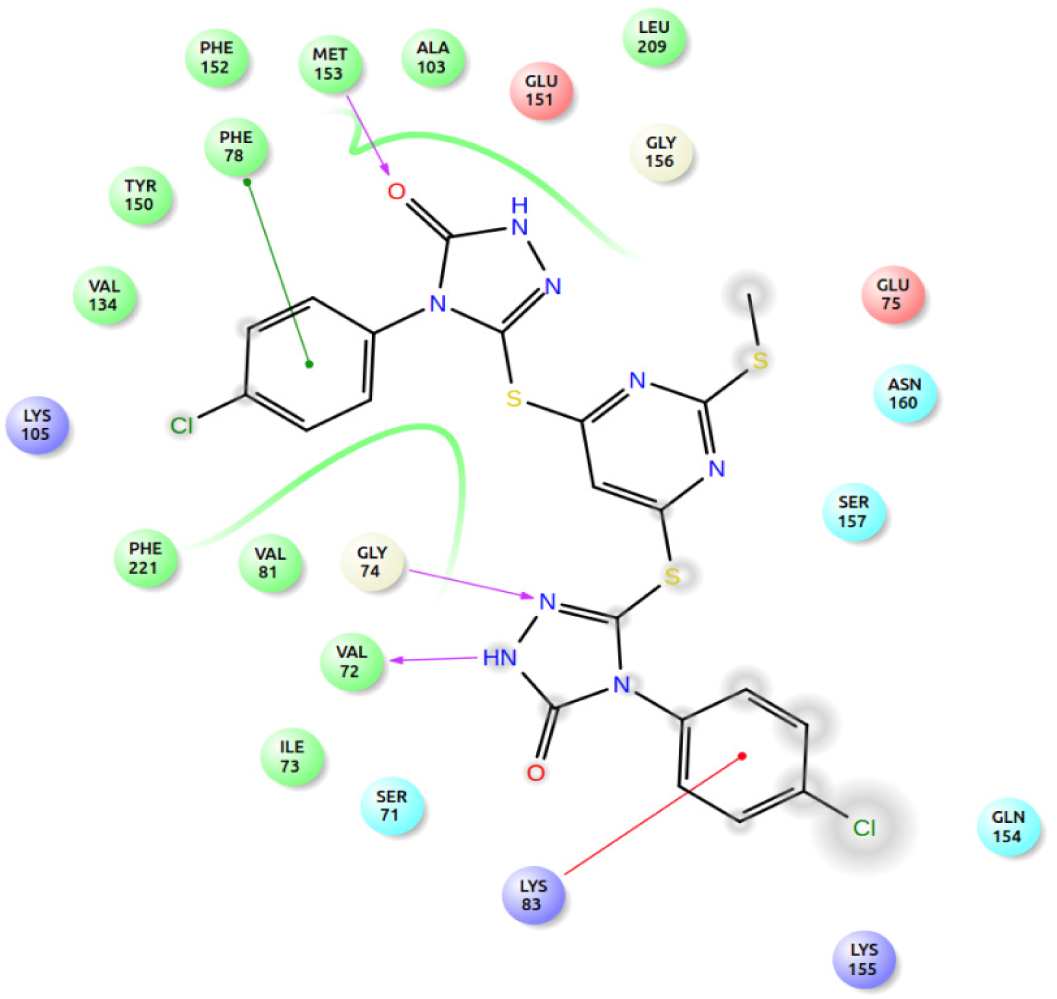
Ligand interaction diagram: SPB07243 and modeled BIK-1.

The position of the docked SPB07243 within the protein can be viewed in the structural representation (Fig 11). Thus, molecular docking studies suggest that the molecules will more likely interact with the binding site of the BIK-1 and acts as possible antagonist to undergo desired mechanism.

**Figure 11:**
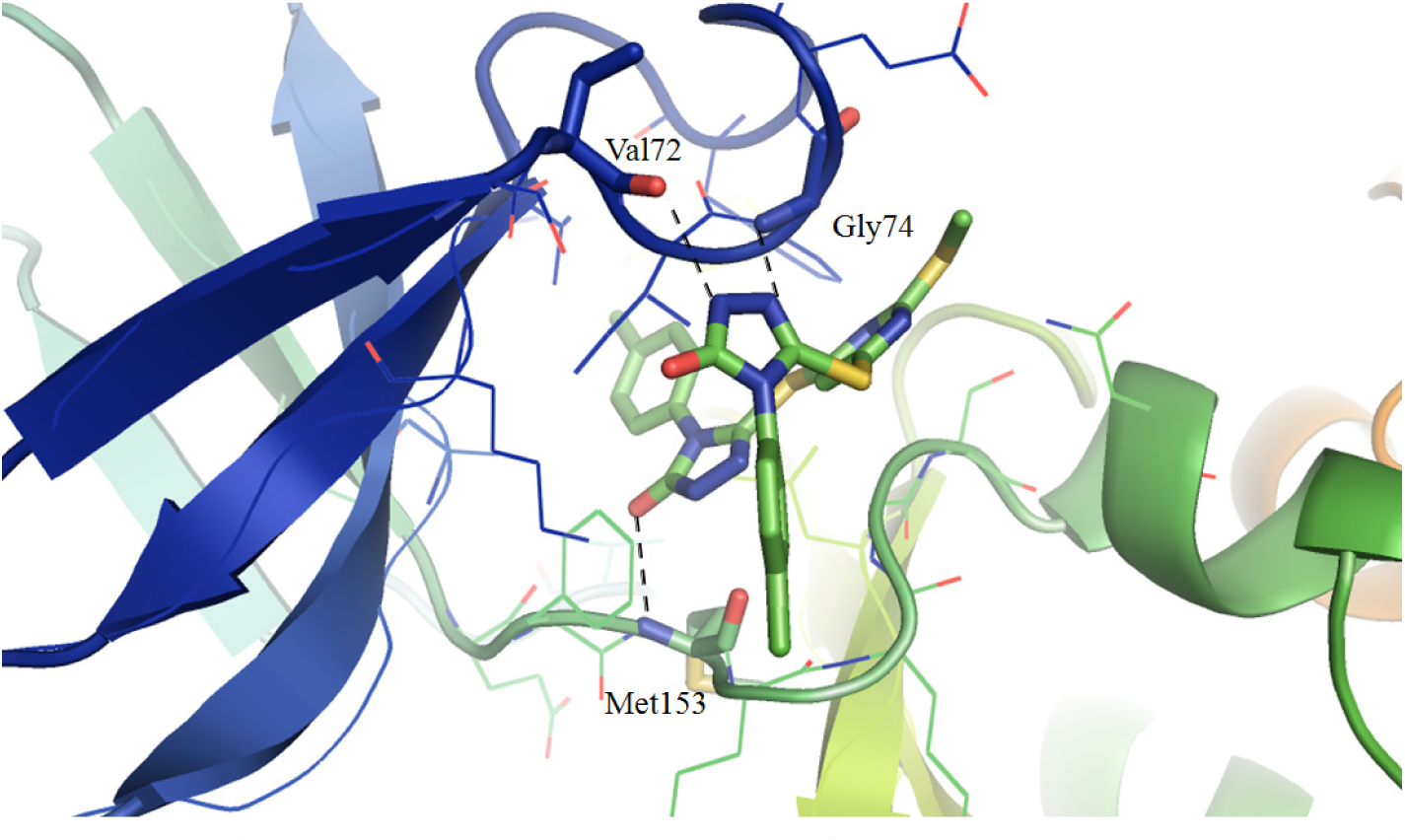
Comprehensive detail of interactions between BIK-1 and SPB07243. Black lines represent hydrogen bonds.

### Molecular Dynamic simulation

To further investigate the stability of the docked complex, 20 ns long MD simulations were carried out for each complex. Each 20 ns long MD simulation run provided a vital confirmation to the docking studies. The energy graph (Fig 12), illustrates the association and dissociation of the ligand molecule with the receptor during the course of simulation. For RF03436 (black), the molecule was adjusting itself into the docked site throughout the simulation. It implies that the molecule, RF03436 was not able to adjust itself in the docked conformation and hence there were interaction energy variation throughout the simulation. However, for SPB07243 (red), the interaction energy was found rather stable (Fig 12). There was sudden drift in the interaction energy at two instances (i.e. at 4 ns and 16 ns), both the time, the -50 KJ/mol drift was noticed. It is estimate that the ligand found better position during the course of simulation. It was vital to identify the primary residues of the BIK-1, which primarily involved in the hydrogen bonding with ligands during the 20 ns long MD simulation. Higher number of hydrogen bonds corresponds to higher strength of the protein ligand complex and hence, its stability. Therefore, VMD analysis was carried out.

**Figure 12:**
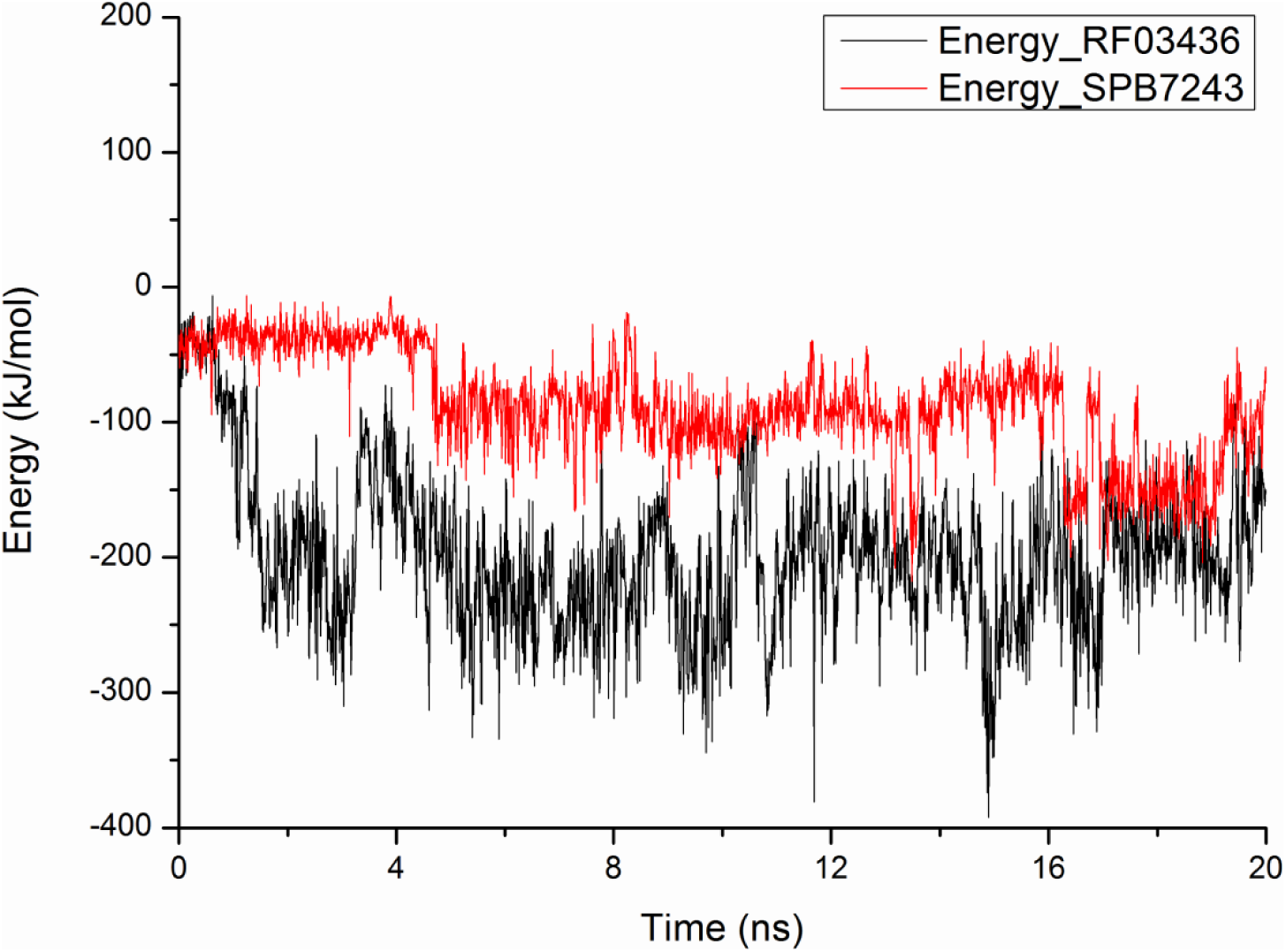
Variation in the interaction energy through 20 ns long MD run.

The analysis revealed that SPB07243 is better molecule for BIK-1 as compared to RF03436. For, RF03436, maximum contribution of hydrogen bonds were made by Tyr214 (23.88%) and Lys155 (11.34%), as evident from the Fig 13A. Among these two residues, only Lys155 is the constituent of the binding site of the protein. In contrast to this, SPB07243 makes hydrogen bonds primarily with Ser219 (48.25%) and Asn207 (12.49%), as evident from Fig 13B.

**Figure 13:**
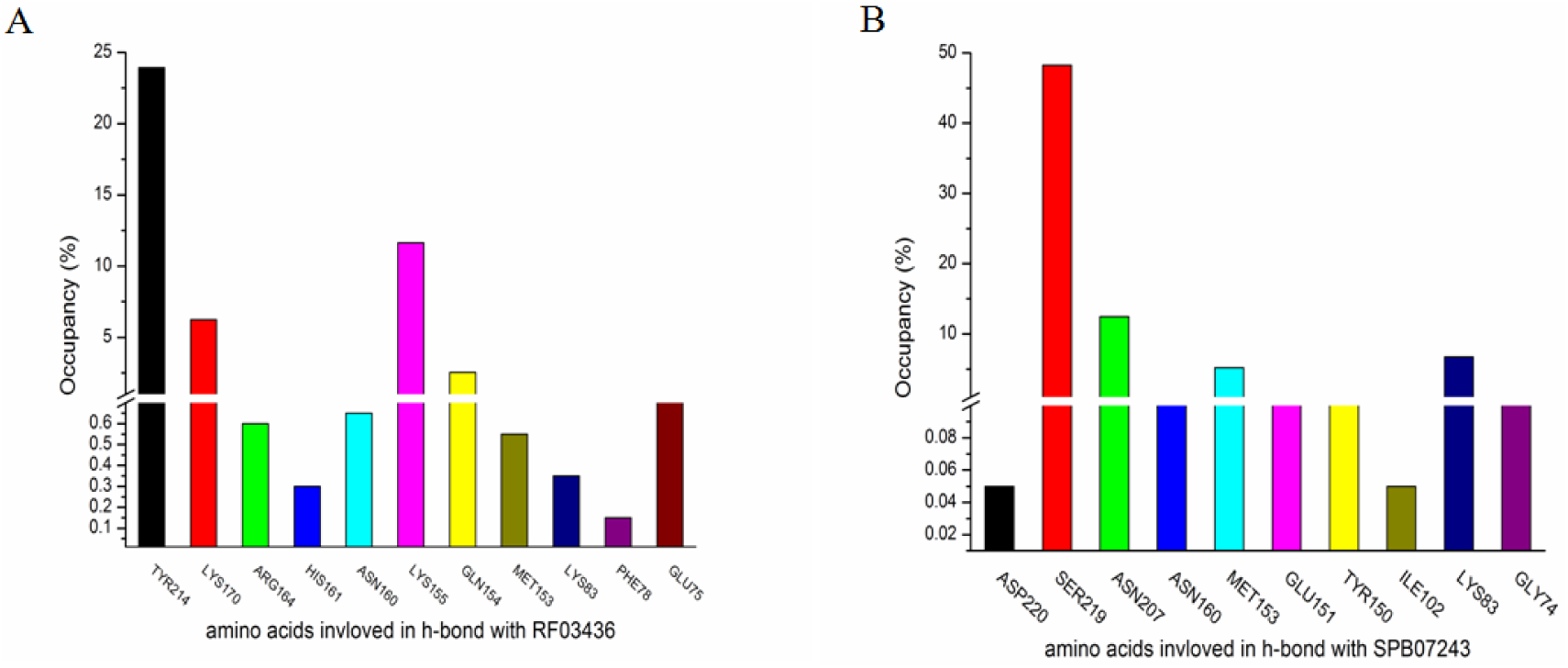
VMD analysis: Hydrogen bond occupancy.

Both the residues are constituent of the binding site of BIK-1. RMSD of the ligand from its initial position was also evaluated. For both the ligands, the RMSD stabilizes at 8 ns, confirming that both the ligand has found better position in the binding site (Fig 14).

**Figure 14:**
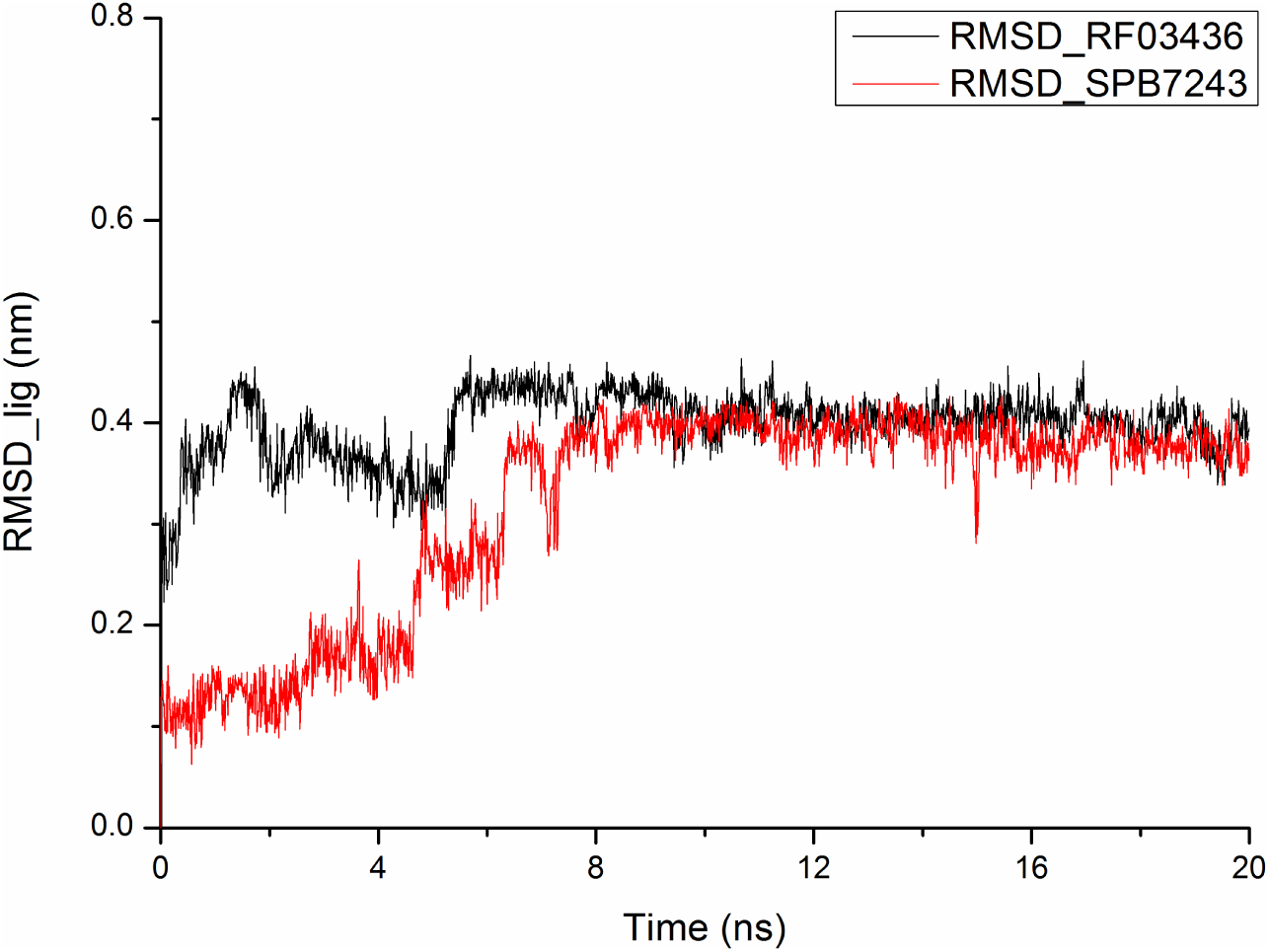
RMSD of ligand with reference to its initial position, during 20ns simulation.

Comparatively, it was found that the RMSD was least for SPB07243. Thus, it can be concluded that SPB07243 could act as better antagonist for BIK-1. Hence, it may be used as bio pesticide against several hemipterans responsible for a huge amount of annual crop loss and significantly increase the rate of plant biomass production.

## Conclusion

We studied through a proposed extension of plant-insect interaction network about how their molecular counterparts of plant immune system interact as well as significantly control the ecological response. We studied the behavior of insect infestation on plants and how these are regulated by PAD4 concentration. Our study based on mathematical model and simulation strongly supports the experimental evidences claiming BIK1 based plant quality control. Based on experimental claims, our model’s further extension also helped studying effect of BIK1 inhibition through *In*_*BIK1*_ in plants resulting as increased plant biomass production. We successfully showed how at the introduction of an inhibitor to BIK1 may help this system to bring down BIK1 activity and influence the PAD4 action. Due to physical limitations of real life scenario, we had to restrict the minimum BIK1 concentration to zero as it reached negative values for some instants and promoted PAD4 activity exponentially high. Results of this model simulation with inhibitor showed how we can enhance crop production by inhibitor based strategies of crop protection from insect infestation.

As an applied part of our proposed model, our work also included the Search for possible inhibitors against BIK1. Out of 53,000 molecules from Maybridge library, there are 88 molecules which can possibly participate in the inhibition mechanisms of BIK-1. However, RF03436 and SPB07243 from the list of 88 possible BIK-1 inhibitors are suggested to be most efficient on the basis of their molecular interactions with the protein. The molecular dynamics simulation studies carried out for the two molecules bound to the protein advocates the higher stability of the BIK-1 and SPB07243 complex. To conclude, it can be fairly assumed that the molecules SPB07243 and RF03436 could be useful for crop protection and improvement policies. Evenmore, SPB07243 seems to be a better option to start with. As well as, our model extension is a contribution to the field of mathematical modeling of Plant-Insect interaction system and their dependencies on their own molecular counterparts.

## Supplement Materials

**Table S1:**
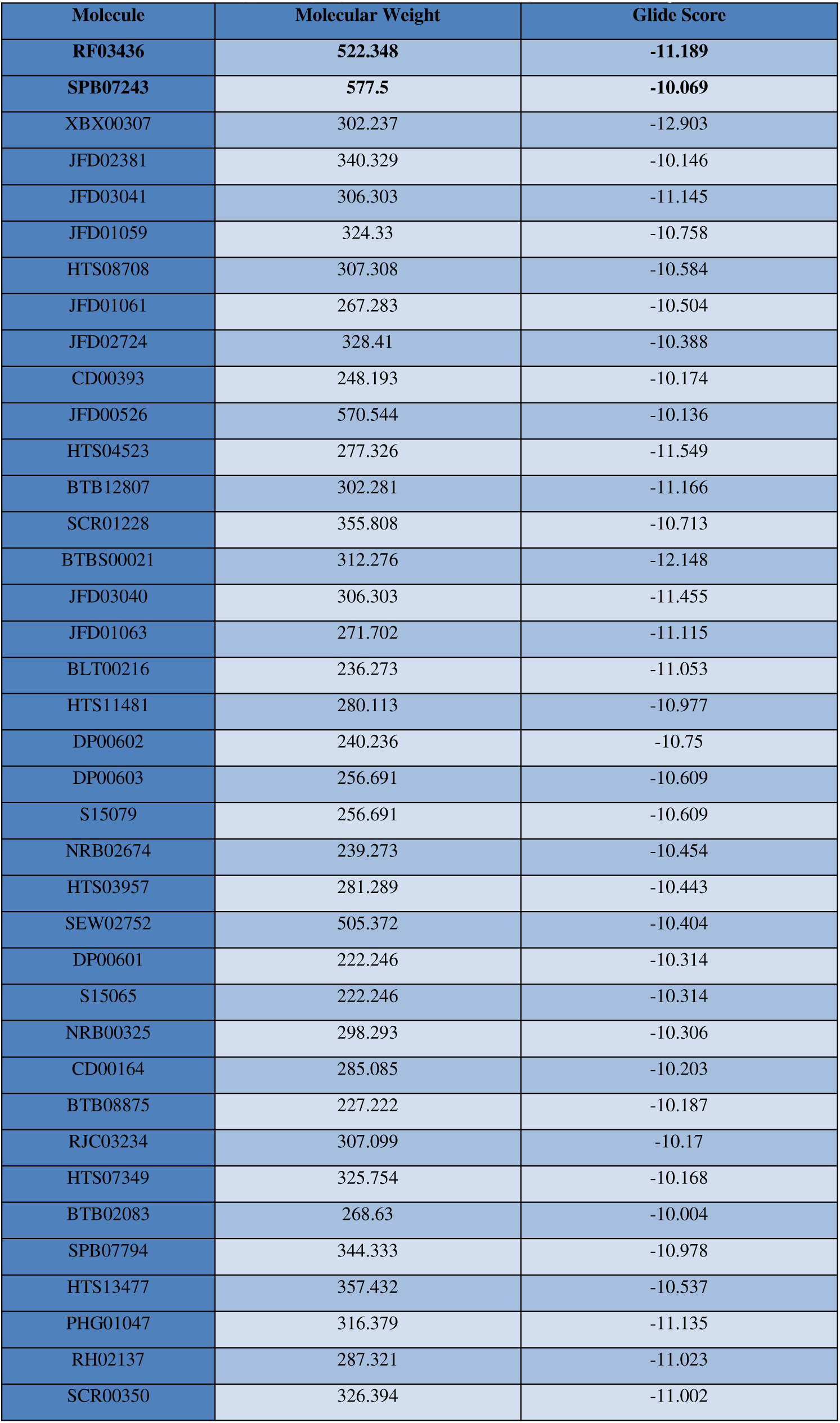
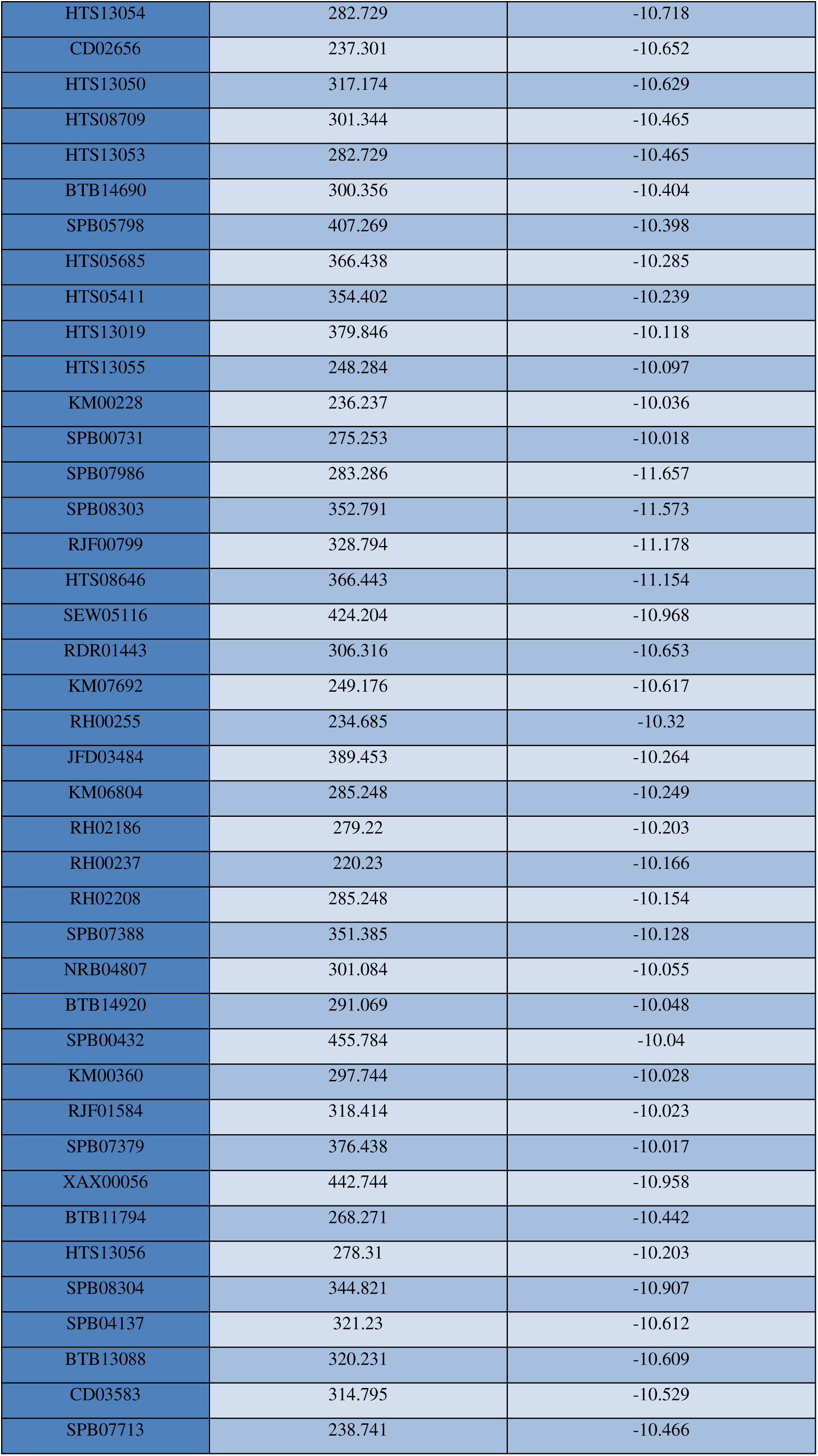
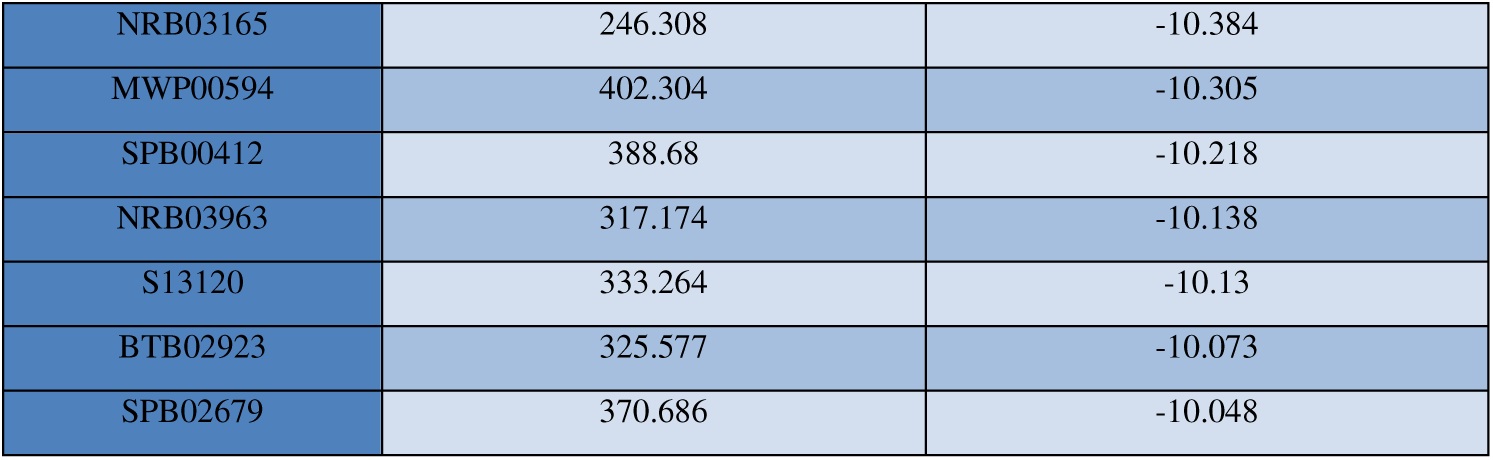
List of all 88 top scoring ligands found interacting with the BIK1 protein.

| Molecule | Molecular Weight | Glide Score |
| --- | --- | --- |
| **RF03436** | **522.348** | **-11.189** |
| **SPB07243** | **577.5** | **-10.069** |
| XBX00307 | 302.237 | -12.903 |
| JFD02381 | 340.329 | -10.146 |
| JFD03041 | 306.303 | -11.145 |
| JFD01059 | 324.33 | -10.758 |
| HTS08708 | 307.308 | -10.584 |
| JFD01061 | 267.283 | -10.504 |
| JFD02724 | 328.41 | -10.388 |
| CD00393 | 248.193 | -10.174 |
| JFD00526 | 570.544 | -10.136 |
| HTS04523 | 277.326 | -11.549 |
| BTB12807 | 302.281 | -11.166 |
| SCR01228 | 355.808 | -10.713 |
| BTBS00021 | 312.276 | -12.148 |
| JFD03040 | 306.303 | -11.455 |
| JFD01063 | 271.702 | -11.115 |
| BLT00216 | 236.273 | -11.053 |
| HTS11481 | 280.113 | -10.977 |
| DP00602 | 240.236 | -10.75 |
| DP00603 | 256.691 | -10.609 |
| S15079 | 256.691 | -10.609 |
| NRB02674 | 239.273 | -10.454 |
| HTS03957 | 281.289 | -10.443 |
| SEW02752 | 505.372 | -10.404 |
| DP00601 | 222.246 | -10.314 |
| S15065 | 222.246 | -10.314 |
| NRB00325 | 298.293 | -10.306 |
| CD00164 | 285.085 | -10.203 |
| BTB08875 | 227.222 | -10.187 |
| RJC03234 | 307.099 | -10.17 |
| HTS07349 | 325.754 | -10.168 |
| BTB02083 | 268.63 | -10.004 |
| SPB07794 | 344.333 | -10.978 |
| HTS13477 | 357.432 | -10.537 |
| PHG01047 | 316.379 | -11.135 |
| RH02137 | 287.321 | -11.023 |
| SCR00350 | 326.394 | -11.002 |
| HTS13054 | 282.729 | -10.718 |
| CD02656 | 237.301 | -10.652 |
| HTS13050 | 317.174 | -10.629 |
| HTS08709 | 301.344 | -10.465 |
| HTS13053 | 282.729 | -10.465 |
| BTB14690 | 300.356 | -10.404 |
| SPB05798 | 407.269 | -10.398 |
| HTS05685 | 366.438 | -10.285 |
| HTS05411 | 354.402 | -10.239 |
| HTS13019 | 379.846 | -10.118 |
| HTS13055 | 248.284 | -10.097 |
| KM00228 | 236.237 | -10.036 |
| SPB00731 | 275.253 | -10.018 |
| SPB07986 | 283.286 | -11.657 |
| SPB08303 | 352.791 | -11.573 |
| RJF00799 | 328.794 | -11.178 |
| HTS08646 | 366.443 | -11.154 |
| SEW05116 | 424.204 | -10.968 |
| RDR01443 | 306.316 | -10.653 |
| KM07692 | 249.176 | -10.617 |
| RH00255 | 234.685 | -10.32 |
| JFD03484 | 389.453 | -10.264 |
| KM06804 | 285.248 | -10.249 |
| RH02186 | 279.22 | -10.203 |
| RH00237 | 220.23 | -10.166 |
| RH02208 | 285.248 | -10.154 |
| SPB07388 | 351.385 | -10.128 |
| NRB04807 | 301.084 | -10.055 |
| BTB14920 | 291.069 | -10.048 |
| SPB00432 | 455.784 | -10.04 |
| KM00360 | 297.744 | -10.028 |
| RJF01584 | 318.414 | -10.023 |
| SPB07379 | 376.438 | -10.017 |
| XAX00056 | 442.744 | -10.958 |
| BTB11794 | 268.271 | -10.442 |
| HTS13056 | 278.31 | -10.203 |
| SPB08304 | 344.821 | -10.907 |
| SPB04137 | 321.23 | -10.612 |
| BTB13088 | 320.231 | -10.609 |
| CD03583 | 314.795 | -10.529 |
| SPB07713 | 238.741 | -10.466 |
| NRB03165 | 246.308 | -10.384 |
| MWP00594 | 402.304 | -10.305 |
| SPB00412 | 388.68 | -10.218 |
| NRB03963 | 317.174 | -10.138 |
| S13120 | 333.264 | -10.13 |
| BTB02923 | 325.577 | -10.073 |
| SPB02679 | 370.686 | -10.048 |

## References

1. Girousse, C. Aphid Infestation Causes Different Changes in Carbon and Nitrogen Allocation in Alfalfa Stems as Well as Different Inhibitions of Longitudinal and Radial Expansion. PLANT Physiol. 137, 1474–1484 (2005).

2. Bos, J. I. B. et al.. A functional genomics approach identifies candidate effectors from the aphid species Myzus persicae (green peach aphid). PLoS Genet. 6, e1001216 (2010).

3. De Vos, M. & Jander, G. Myzus persicae (green peach aphid) salivary components induce defence responses in Arabidopsis thaliana. Plant Cell Environ. 32, 1548–1560 (2009).

4. Will, T. & van Bel, A. J. E. Physical and chemical interactions between aphids and plants. J. Exp. Bot. 57, 729–737 (2006).

5. Tjallingii, W. F. Salivary secretions by aphids interacting with proteins of phloem wound responses. J. Exp. Bot. 57, 739–745 (2006).

6. Goggin, F. L. Plant-aphid interactions: molecular and ecological perspectives. Curr. Opin. Plant Biol. 10, 399–408 (2007).

7. Control Theory and Systems Biology. (The MIT Press, 2009).

8. Edelstein-Keshet, L. Mathematical theory for plant—herbivore systems. J. Math. Biol. 24, 25–58 (1986).

9. Stieha, C. R., Abbott, K. C. & Poveda, K. The Effects of Plant Compensatory Regrowth and Induced Resistance on Herbivore Population Dynamics. Am. Nat. 187, 167–181 (2016).

10. Kartal, S. Dynamics of a plant herbivore model with differential difference equations. Cogent Math. 3, (2016).

11. May, R. M. *Stability and complexity in model ecosystems*. (Princeton University Press, 2001).

12. Chattopadhayay, J., Sarkar, R., Fritzsche-Hoballah, M. E., Turlings, T. C. J. & Bersier, L.-F. Parasitoids may Determine Plant Fitness—A Mathematical Model Based on Experimental Data. J. Theor. Biol. 212, 295–302 (2001).

13. Danca, M., Codreanu, S. & Bakó, B. Detailed analysis of a nonlinear prey-predator model. J. Biol. Phys. 23, 11–20 (1997).

14. Lei, J., A Finlayson, S., Salzman, R. A., Shan, L. & Zhu-Salzman, K. BOTRYTIS-INDUCED KINASE1 Modulates Arabidopsis Resistance to Green Peach Aphids via PHYTOALEXIN DEFICIENT4. Plant Physiol. 165, 1657–1670 (2014).

15. Mantelin, S., Bhattarai, K. K. & Kaloshian, I. Ethylene contributes to potato aphid susceptibility in a compatible tomato host. New Phytol. 183, 444–456 (2009).

16. Thompson, G. A. & Goggin, F. L. Transcriptomics and functional genomics of plant defence induction by phloem-feeding insects. J. Exp. Bot. 57, 755–766 (2006).

17. Kettles, G. J., Drurey, C., Schoonbeek, H., Maule, A. J. & Hogenhout, S. A. Resistance of Arabidopsis thaliana to the green peach aphid, Myzus persicae, involves camalexin and is regulated by microRNAs. New Phytol. 198, 1178–1190 (2013).

18. Jirage, D. et al.. Arabidopsis thaliana PAD4 encodes a lipase-like gene that is important for salicylic acid signaling. Proc. Natl. Acad. Sci. U. S. A. 96, 13583–13588 (1999).

19. Pegadaraju, V. Premature Leaf Senescence Modulated by the Arabidopsis PHYTOALEXIN DEFICIENT4 Gene Is Associated with Defense against the Phloem-Feeding Green Peach Aphid. PLANT Physiol. 139, 1927–1934 (2005).

20. Pegadaraju, V. et al.. Phloem-based resistance to green peach aphid is controlled by Arabidopsis PHYTOALEXIN DEFICIENT4 without its signaling partner ENHANCED DISEASE SUSCEPTIBILITY1. Plant J. Cell Mol. Biol. 52, 332–341 (2007).

21. Louis, J. et al.. Discrimination of Arabidopsis PAD4 activities in defense against green peach aphid and pathogens. Plant Physiol. 158, 1860–1872 (2012).

22. Louis, J. & Shah, J. Plant defence against aphids: the PAD4 signalling nexus. J. Exp. Bot. 66, 449–454 (2015).

23. Louis, J., Leung, Q., Pegadaraju, V., Reese, J. & Shah, J. *PAD4* -Dependent Antibiosis Contributes to the *ssi2* -Conferred Hyper-Resistance to the Green Peach Aphid. Mol. Plant. Microbe Interact. 23, 618–627 (2010).

24. Lu, D. et al.. A receptor-like cytoplasmic kinase, BIK1, associates with a flagellin receptor complex to initiate plant innate immunity. Proc. Natl. Acad. Sci. 107, 496–501 (2010).

25. Zhang, J. et al.. Receptor-like Cytoplasmic Kinases Integrate Signaling from Multiple Plant Immune Receptors and Are Targeted by a Pseudomonas syringae Effector. Cell Host Microbe 7, 290–301 (2010).

26. Bent, A. F. & Mackey, D. Elicitors, effectors, and R genes: the new paradigm and a lifetime supply of questions. Annu. Rev. Phytopathol. 45, 399–436 (2007).

27. Boller, T. & Felix, G. A renaissance of elicitors: perception of microbe-associated molecular patterns and danger signals by pattern-recognition receptors. Annu. Rev. Plant Biol. 60, 379–406 (2009).

28. Prince, D. C., Drurey, C., Zipfel, C. & Hogenhout, S. A. The leucine-rich repeat receptor-like kinase BRASSINOSTEROID INSENSITIVE1-ASSOCIATED KINASE1 and the cytochrome P450 PHYTOALEXIN DEFICIENT3 contribute to innate immunity to aphids in Arabidopsis. Plant Physiol. 164, 2207–2219 (2014).

29. Evans, D. J. A new 4th order runge-kutta method for initial value problems with error control. Int. J. Comput. Math. 39, 217–227 (1991).

30. Ferreira, L., dos Santos, R., Oliva, G. & Andricopulo, A. Molecular Docking and Structure-Based Drug Design Strategies. Molecules 20, 13384–13421 (2015).

31. Nguyen, G. T. T. et al.. Chalcone-based Selective Inhibitors of a C4 Plant Key Enzyme as Novel Potential Herbicides. Sci. Reports 6, (2016).

32. Combet, C., Blanchet, C., Geourjon, C. & Deléage, G. NPS@: network protein sequence analysis. Trends Biochem. Sci. 25, 147–150 (2000).

33. Altschul, S. F. et al.. Gapped BLAST and PSI-BLAST: a new generation of protein database search programs. Nucleic Acids Res. 25, 3389–3402 (1997).

34. Berman, H. M. The Protein Data Bank. Nucleic Acids Res. 28, 235–242 (2000).

35. Fiser, A. in Computational Biology (ed. Fenyö, D.) 673, 73–94 (Humana Press, 2010).

36. Clore, G. M., Robien, M. A. & Gronenborn, A. M. Exploring the limits of precision and accuracy of protein structures determined by nuclear magnetic resonance spectroscopy. J. Mol. Biol. 231, 82–102 (1993).

37. Hardin, C., Pogorelov, T. V. & Luthey-Schulten, Z. Ab initio protein structure prediction. Curr. Opin. Struct. Biol. 12, 176–181 (2002).

38. Kelley, L. A., Mezulis, S., Yates, C. M., Wass, M. N. & Sternberg, M. J. E. The Phyre2 web portal for protein modeling, prediction and analysis. Nat. Protoc. 10, 845–858 (2015).

39. Zhang, J., Liang, Y. & Zhang, Y. Atomic-Level Protein Structure Refinement Using Fragment-Guided Molecular Dynamics Conformation Sampling. Structure 19, 1784–1795 (2011).

40. Lovell, S. C. et al.. Structure validation by Calpha geometry: phi,psi and Cbeta deviation. Proteins 50, 437–450 (2003).

41. Wiederstein, M. & Sippl, M. J. ProSA-web: interactive web service for the recognition of errors in three-dimensional structures of proteins. Nucleic Acids Res. 35, W407–410 (2007).

42. Volkamer, A., Kuhn, D., Grombacher, T., Rippmann, F. & Rarey, M. Combining global and local measures for structure-based druggability predictions. J. Chem. Inf. Model. 52, 360–372 (2012).

43. Zhang, Z., Li, Y., Lin, B., Schroeder, M. & Huang, B. Identification of cavities on protein surface using multiple computational approaches for drug binding site prediction. Bioinforma. Oxf. Engl. 27, 2083–2088 (2011).

44. Huang, B. MetaPocket: a meta approach to improve protein ligand binding site prediction. Omics J. Integr. Biol. 13, 325–330 (2009).

45. Marchler-Bauer, A. et al.. CDD/SPARCLE: functional classification of proteins via subfamily domain architectures. Nucleic Acids Res. 45, D200–D203 (2017).

46. Sigrist, C. J. A. et al.. PROSITE, a protein domain database for functional characterization and annotation. Nucleic Acids Res. 38, D161–D166 (2010).

47. Xu, J. et al.. Identification and functional analysis of phosphorylation residues of the Arabidopsis BOTRYTIS-INDUCED KINASE1. Protein Cell 4, 771–781 (2013).

48. Letunic, I., Doerks, T. & Bork, P. SMART: recent updates, new developments and status in 2015. Nucleic Acids Res. 43, D257–D260 (2015).

49. Jorgensen, W. L., Maxwell, D. S. & Tirado-Rives, J. Development and Testing of the OPLS All-Atom Force Field on Conformational Energetics and Properties of Organic Liquids. J. Am. Chem. Soc. 118, 11225–11236 (1996).

50. Berendsen, H. J. C., van der Spoel, D. & van Drunen, R. GROMACS: A message-passing parallel molecular dynamics implementation. Comput. Phys. Commun. 91, 43–56 (1995).

51. Scott, W. R. P. et al.. The GROMOS Biomolecular Simulation Program Package. J. Phys. Chem. A 103, 3596–3607 (1999).

52. Schüttelkopf, A. W. & van Aalten, D. M. F. PRODRG: a tool for high-throughput crystallography of protein-ligand complexes. Acta Crystallogr. D Biol. Crystallogr. 60, 1355–1363 (2004).

53. Humphrey, W., Dalke, A. & Schulten, K. VMD: visual molecular dynamics. J. Mol. Graph. 14, 33–38, 27–28 (1996).

